# Intelectin-2 is a broad-spectrum antimicrobial lectin

**DOI:** 10.1101/2025.06.09.658748

**Authors:** Amanda E. Dugan, Deepsing Syangtan, Eric B. Nonnecke, Rajeev S. Chorghade, Amanda L. Peiffer, Jenny J. Yao, Jessie Ille-Bunn, Dallis Sergio, Gleb Pishchany, Catherine Dhennezel, Hera Vlamakis, Sunhee Bae, Sheila Johnson, Chariesse Ellis, Soumi Ghosh, Jill W. Alty, Carolyn E. Barnes, Miri Krupkin, Gerardo Cárcamo-Oyarce, Katharina Ribbeck, Ramnik J. Xavier, Charles L. Bevins, Laura L. Kiessling

## Abstract

Mammals regulate the localization, composition, and activity of their native microbiota at colonization sites. Lectins residing at these sites influence microbial populations, but their individual functions are often unclear. Intelectins are found in chordates at mucosal barriers, but their functions are not well characterized. We found that mouse intelectin-2 (mItln2) and human intelectin-2 (hItln2) engage and crosslink mucins via carbohydrate recognition. Moreover, both lectins recognize microbes within native microbial communities, including gram-positive and gram-negative isolates from the respiratory and gastrointestinal tracts. This ability to engage mammalian and microbial glycans arises from calcium-coordinated binding of carbohydrate residues within mucus and microbial surfaces. Microbes, but not human cells, bound by mItln2 or hItln2, suffer a loss of viability. These findings underscore the crucial antimicrobial role of mammalian intelectin-2 in mucosal defense, where it plays offensive (microbial killing) and defensive (mucus crosslinking) roles in regulating microbial colonization.

## INTRODUCTION

Mucosal surfaces of the mammalian respiratory and gastrointestinal tracts are replete with microorganisms seeking to colonize these host tissues. Under homeostatic conditions, microbial communities flourish and contribute to host health. Simultaneously, exposure to opportunistic pathogens, allergens, and helminths can injure host tissues and lead to infection. Host tissues must therefore respond to an ever-changing landscape of immunological challenges.

In response to microbial colonization, host cells secrete a suite of soluble factors that regulate the localization, composition, and activity of microbial communities. Among these factors are lectins, carbohydrate-binding proteins that bind glycans on host and microbial cells^1–4^. The intelectins, or X-type lectins, are secreted at mucosal surfaces and, though purported to play a role in host defense against microbes, aspects of their function are still unclear^5–11^. Human and mouse intelectin-1 (Itln1) are constitutively expressed in the intestines and exclusively recognize a set of microbial glycans^12–16^. Itln1 has been suggested to facilitate phagocytic uptake of microbes by neutrophils and regulate the localization of mucolytic microbes^8,10,13^. In addition to Itln1, humans and certain mouse strains possess a second intelectin gene, intelectin-2 (Itln2), which has poorly characterized ligand specificity and unclear biological function^6,17–20^. Mouse intelectin-2 (mItln2) has been reported to be upregulated during nematode infections. Human intelectin-2 (ITLN2, henceforth hItln2) is constitutively expressed and restricted to the small intestine. Moreover, in the small intestine of ileal Crohn’s disease (CD) patients, the expression of hItln2 is decreased while it is increased in the colonic tissue of patients with colonic CD and ulcerative colitis (UC). Given that intelectin-2 is implicated in inflammatory processes and disorders involving microbial dysbiosis, we investigated its ligand specificity and function^15,21–26^.

Herein, we report the characterization of mItln2 and hItln2. Using a novel mouse model with an intact intelectin locus, we found that mItln2 is selectively and potently induced by T-helper type 2 (Th2) stimuli. We show that mItln2 and hItln2 are indeed lectins, as they recognize select glycans with terminal β-D-galactopyranose residues, which are found on mammalian and microbial glycans. Regarding the host, these lectins crosslink mucins in a glycan-dependent manner. Regarding microbes, these lectins bind a range of gram-positive and gram-negative bacteria. Moreover, we found that mItln2 exerts direct microbicidal effects. HItln2 binding also limits microbial growth, in this case through agglutination, and its activity is intriguingly more potent at lower pH and salt conditions. Taken together, our findings reveal the roles of Itln2 in host defense – through mucin crosslinking and broad-spectrum antimicrobial activity–underscoring the multifaceted roles of intelectins in host–microbe interactions.

## RESULTS

### MItln2 expression is induced by IL-4 and IL-13 in enteroids

In Th2 inflammation, microbial-, helminth-, and allergen-derived toxins and proteases compromise the integrity of host mucosal barriers^27–29^. Such damage can increase the risk of infection by opportunistic pathobionts and prompt the host to activate wound repair and defense pathways. There is evidence that mammalian intelectins can be induced during Th2-type immune responses, including during intestinal helminth infections and asthma, but many details are unclear^6,7,17,20,23,24,30^.

We used a mouse model to probe the regulation of intelectin expression during inflammation. The widely used strains for studying inflammation, such as C57BL/6 and its sub-strains (e.g., C57BL/10), encode a single intelectin gene, *Itln1*, resulting from a large 420-kb deletion across the intelectin locus^9,17,18,31,32^. However, most wild-derived and conventional laboratory mouse strains encode six intelectin paralogs: *Itln1*, *Itln2*, *Itln3*, *Itln4*, *Itln5*, and *Itln6*^18,31^. To better understand intelectin regulation, we generated a congenic C57BL/6 mouse model (B6.C-*Itln1-6*) that encodes a complete *Itln1-6* locus derived from the BALB/c strain^18,33^. Stimulation of enteroids from B6.C-*Itln1-6* mice with type 2 cytokines (IL-4 and IL-13) resulted in a six-fold increase in total *Itln* expression compared to a two-fold increase in wild-type C57BL/6 enteroids (Fig. 1a,b). In tissue and unstimulated enteroids from B6.C-*Itln1-6* mice, most (≥ 94%) of total intelectin mRNA was *Itln1,* with 2-5% being *Itln2* (Fig. 1c,d). Following IL-4 and IL-13 treatment, the relative abundance of *Itln2* increased to 30% of total transcript reads, indicating a 40-fold upregulation (Fig. 1d; Supplementary Table 1). This finding aligns with previous report that *Itln2* is induced in the small intestine of BALB/c mice infected with *Nippostrongylus brasiliensis*, *Trichuris muris*, and *Trichinella spiralis*, which are known to promote Th2 inflammation^6,17,20^. No other intelectins (*Itln3-6*) were detected in small intestinal tissue or enteroids at baseline or after type 2 cytokine stimulation. These results establish that Th2-type cytokines selectively induce mItln2 (Fig. 1e), which, at baseline, is expressed at low levels. These data suggest a role of mItln2 distinct from other intelectins.

**Figure 1:**
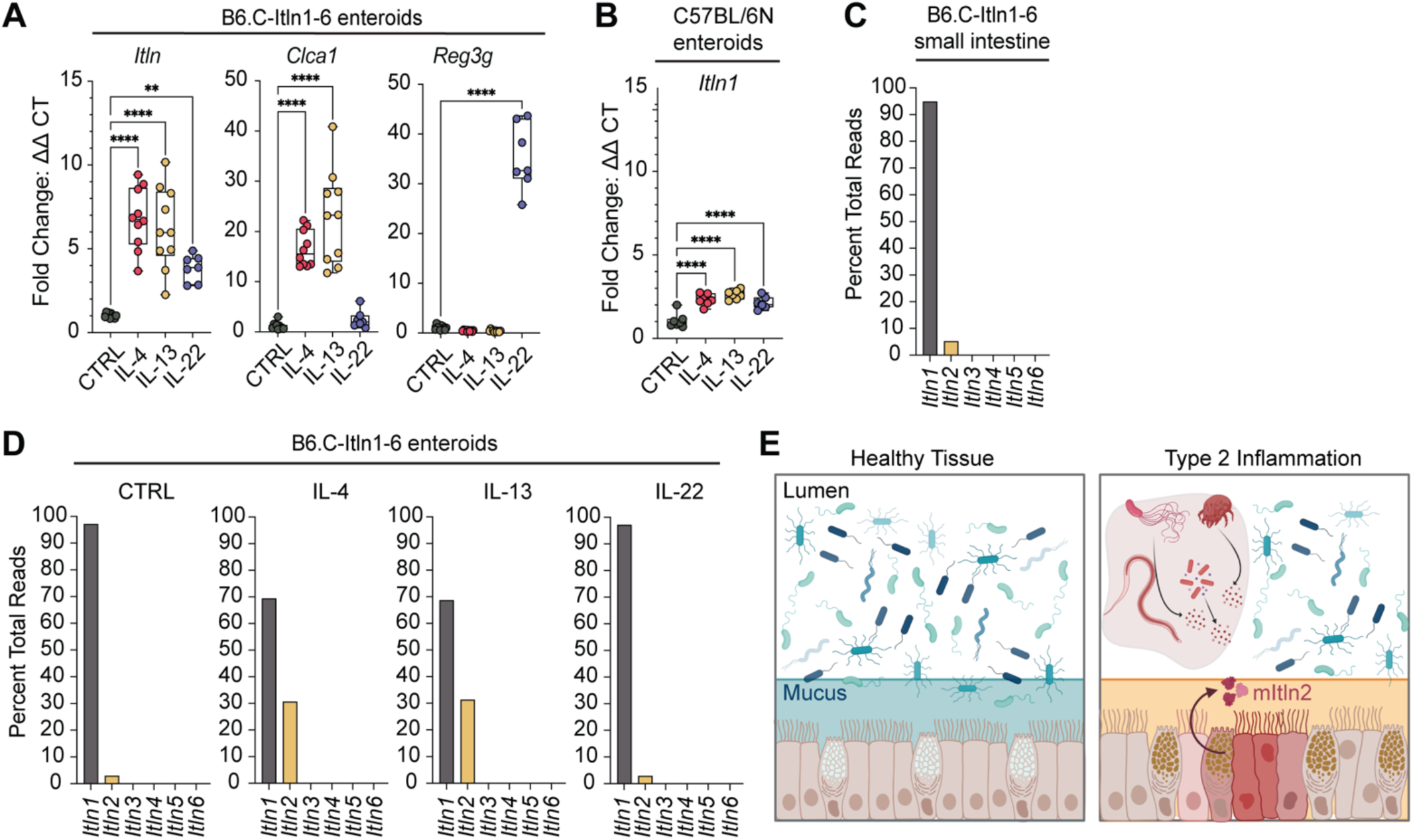
Mouse intelectin-2 (mItln2) is conditionally expressed in type 2 inflammation. **(A)** Relative quantification (mean ± SEM) of total *Itln* expression in enteroids derived from B6.C-*Itln1-6* mice after treatment with different cytokines. Goblet cell product *Clca1* and intestinal epithelial cell product *Reg3g* served as positive controls for IL-4/ IL-13 and IL-22 treatments, respectively (collective data from 3 independent experiments). **(B)** Relative quantification (mean ± SEM) of *Itln1* expression in enteroids derived from C57BL/6N mice after treatment with different cytokines (collective data from 3 independent experiments). **(C** and **D)** Percentage of total *Itln* sequencing reads mapped to different *Itln* paralogs. Data represent percent reads from bulk RNAseq analysis of pooled samples from distal small intestine in B6.C- *Itln1-6* mice (n = 3) (C) and B6.C-*Itln1-6*-derived enteroids stimulated with different cytokines (n = 6 independent experiments) (D). (**E**) Schematic of mItln2 expression in mouse lungs and intestines upon exposure to helminths and allergens. Statistical analysis in 1A and 1B involves one-way ANOVA with Dunnett’s multiple comparisons test. **P < 0.01, ****P < 0.0001.

### MItln2 recognizes select glycans with terminal β-D-galactopyranose residues

mItln2 shares 81% sequence identity with the homologous protein Itln1. This similarity prompted us to analyze the structure and function of mItln2. To this end, we generated a recombinant form of mItln2 containing a StrepII tag that facilitated purification and detection (Supplementary Fig. 1a,b). We found that strep-mItln2 was glycosylated and properly folded with a melting temperature of approximately 53 °C (Supplementary Fig. 1c– h). Circular dichroism analysis further revealed that mItln2 exhibits a mixed alpha-helical and beta-sheet secondary structure, suggesting a comparable folded structure as Itln1 (Supplementary Fig. 1c). Although mItln2 lacks the two cysteines present in hItln1 that form a disulfide-linked homotrimer (Supplementary Fig. 1a)^12,34^, treatment of mItln2 with the bifunctional homo-crosslinker bis-sulfosuccinimidyl suberate gave rise to a laddered mixture of monomer, dimer, trimer, and hexamer species, suggesting non-covalent oligomerization of mItln2 (SupplementarySI Fig. 1i). Dynamic light scattering also showed that mItln2 occupies a variable distribution of oligomeric species in solution (SupplementarySI Fig. 1j). Together, these data indicate that mItln2 is a stably folded, glycosylated lectin that assembles into non-covalent oligomers.

Human and mouse Itln1 were previously shown to have identical binding sites and bind exclusively to microbial carbohydrates, including β-D-galactofuranose (β-Gal*f*), the five-membered ring isomer of galactose^12,14^. Despite the high sequence similarities of mItln2 with mItln1 and hItln1, the ligand-binding domain of mItln2 differs in a single residue (W288A) (Fig. 2a). The aromatic box formed by W288 and Y297 is crucial for Itln1’s specificity for microbial monosaccharides. The parsimonious binding site change between mItln1 and mItln2 suggests that the latter might exhibit different ligand specificities than mItln1 and hItln1, leading to differences in microbial binding^13^.

**Figure 2:**
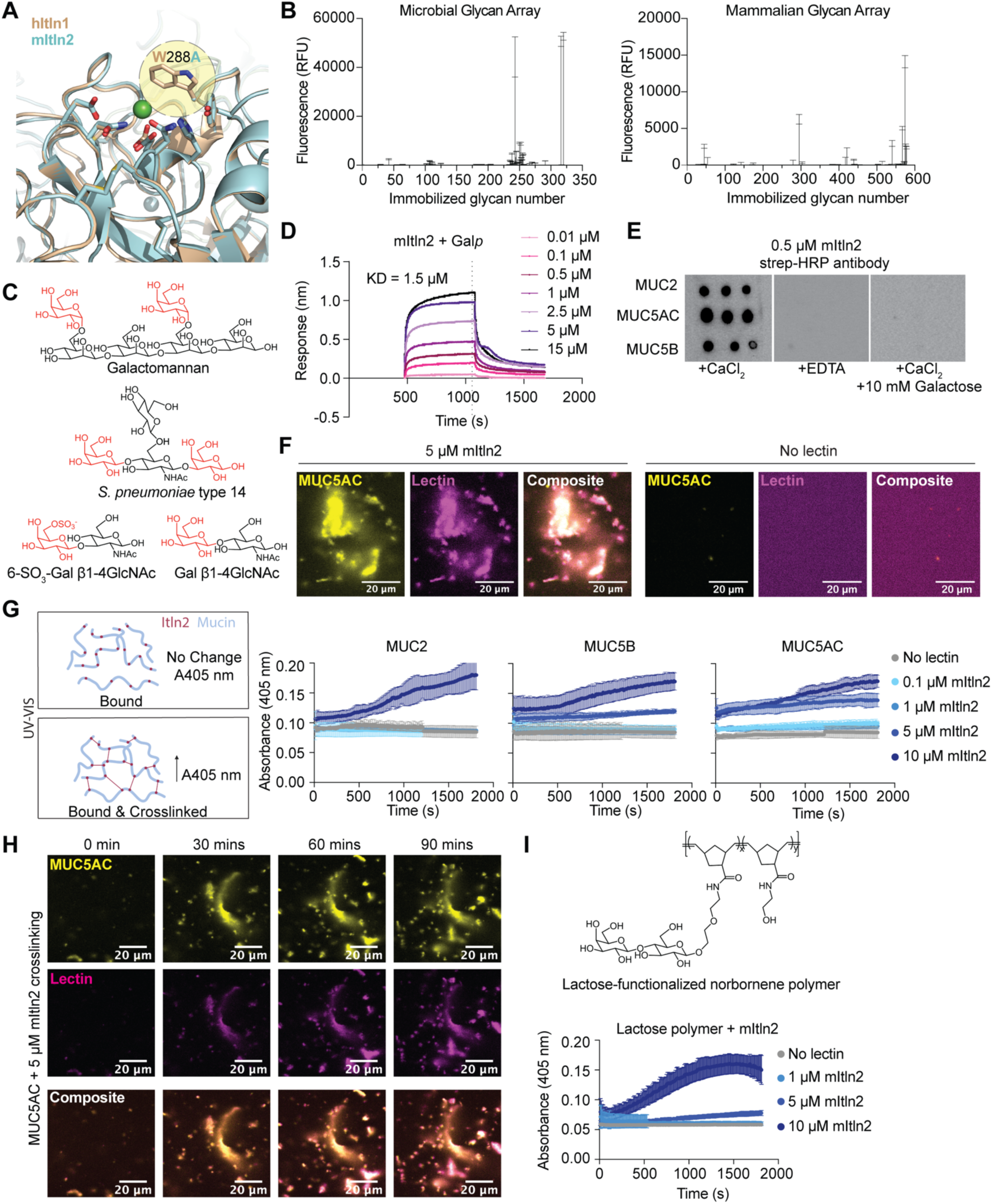
Characterization of glycan specificity for mItln2. **(A)** Overlay of carbohydrate recognition domain of hItln1 (wheat, PDB ID 4WMY) and mItln2 (teal, predicted model), showing calcium (green), conserved calcium coordination residues, and the presence of alanine instead of tryptophan at position 288 in mItln2. hItln1 and mItln1 have identical binding site residues. (B) Binding profiles of recombinant mItln2 (25 µg/mL) to microbial (left) and mammalian (right) glycan microarrays from the National Center for Functional Glycomics (NCFG). Data are shown as mean ± SD (n = 4 technical replicates). RFU, Relative Fluorescence Unit. (**C**) Glycan structures bound by mItln2 in glycan arrays. All hits contain either a terminal Gal*p* or 6SO_3_-Gal*p*, shown in red. (**D**) BLI trace of mItln2 binding to immobilized biotinylated-b-Gal*p* at varying concentrations, with an apparent KD of 1.5 µM. Data were normalized by background subtraction, using biotin-loaded streptavidin. (**E**) Dot blot analysis of MUC2, MUC5AC, and MUC5B (1 μg each) on nitrocellulose membrane, probed with 0.5 μM mItln2 in the presence of Ca^2+^ or EDTA. A control with 100 mM galactose was also included. Binding of mItln2 was detected using a Strep-HRP antibody. (**F**) Images of 5 μM mItln2 (magenta) binding to 0.01% (w/v) fluorescently labeled MUC5AC (yellow). Binding of mItln2 was detected with Strep antibody. MUC5AC without lectin treatment served as a control. Scale bars, 20 µm. (**G**) Spectroscopic assay for crosslinking of 0.05% (w/v) mucins with varying amounts of mItln2, measured by the increase in absorbance at 405 nm. Schematic of the crosslinking assay is shown on the left. Data are shown as mean ± SD (n = 3 technical replicates). (**H**) Time-lapse images of 0.01% (w/v) fluorescently labeled MUC5AC (yellow) treated with 5 μM mItln2 (magenta). Binding of mItln2 was detected with Strep antibody. Scale bars, 20 µm. (**I**) Spectroscopic assay for crosslinking of 0.05% (w/v) lactose-functionalized glycopolymer by mItln2, measured by the increase in absorbance at 405 nm. Data are shown as mean ± SD (n = 3 technical replicates). Structure of the lactose-functionalized trans-poly (norbornene) is shown on the top. Results in (D), (E), and (G) are representative of three independent experiments. Results in (F), (H), and (I) are representative of two independent experiments.

To assess whether mItln2 is a glycan-binding protein, we screened the recombinant protein on mammalian and microbial glycan arrays (Supplementary Fig. 2a). Microbial glycan analysis yielded hits, including yeast and plant-like galactomannan (Davanat) and bacterial galactose-containing capsular polysaccharides (Fig. 2b,c; Supplementary Table 2)^34^. In addition, mItln2 bound mammalian glycans, including oligosaccharides from N- and O- glycans displaying terminal 6-SO_3_-galactopyranose or α- or β-D-galactopyranose (Fig. 2b,c; Supplementary Fig. 2b,c; Supplementary Tables 2). Notably, the binding profile of mItln2 had no overlap with hItln1, indicating distinct specificity. We previously reported that hItln1, like mItln1, binds to acyclic vicinal (1,2)-diol moieties present on microbial but not mammalian glycans (Supplementary Fig. 2d)^2,12^.

To quantify the array results and assess mItln2 minimal binding groups, we used an enzyme-linked lectin assay (ELLA). In this assay, biotinylated carbohydrates were immobilized on streptavidin-coated plates and increasing concentrations of mItln2 were added. Plates were washed to remove unbound lectin and bound mItln2 was detected with anti-StrepII-HRP antibody (Supplementary Fig. 2e,f). Consistent with the array results, these experiments demonstrated that mItln2 binds β-*N*-acetyllactosamine (β-LacNAc), and β-6-SO_3_-*N-* acetyllactosamine (β-6SO_3_ LacNAc) (Supplementary Fig. 2g). Because β-LacNAc is a disaccharide composed of terminal β-D-galactopyranose (β-Gal*p*) linked to β-D*-N*-acetyl-glucosamine (β-GlcNAc), we also evaluated binding to individual monosaccharides to determine mItln2 specificity. ELLA results show that mItln2 does not bind β-GlcNAc and only recognizes β-Gal*p* (Supplementary Fig. 2g). The lectin binding was dose-dependent, a hallmark of specific recognition. Additional experiments using biolayer interferometry (BLI) indicate that mItln2 selectively binds the pyranose form of galactose (β-Gal*p*) but not the furanose form (β-Gal*f*) recognized by Itln1 (Supplementary Fig. 2h–j). The apparent dissociation constant of mItln2 for β-Gal*p* is approximately 1.5 μM (Fig. 2d; Supplementary Fig. 2k). Because mItln2 appears to accommodate both β-Gal*p* and β-6SO_3_- Gal*p* as ligands, we postulate that it uses calcium to coordinate binding to conserved hydroxyl groups on the pyranose ring of galactose, a feature shared by all glycan hits. Still, mItln2 does not bind every β-Gal*p-* containing glycan on the array, indicating there may be additional subsites that contribute to affinity. Notably, a single amino acid difference in the binding site of mItln2 allows this lectin to preferentially bind the pyranose form of galactose over the furanose form preferred by Itln1.

### MItln2 crosslinks mucin glycoproteins

During parasitic nematode infections, mucins are secreted by goblet cells into the intestinal and lung mucosa, where they can interact with newly expressed mItln2^17^. Mucins are glycosylated with epitopes similar to those of the top mammalian glycan array hits for mItln2; therefore, we assayed its interaction with gastric and intestinal mucins. We first immobilized commercially available gastric mucus to nitrocellulose in a dot blot assay. Following incubation with mItln2, we observed robust binding to gastric mucus in the presence of calcium. EDTA addition abrogated binding, as expected, given that mItln2 and hItln1 are related members of the calcium-dependent X-type lectin family. These data indicate that mItln2 binds mucus by engaging mucin glycans (Supplementary Fig. 2l). To test glycan binding more directly, we added 10 mM galactose and found it could competitively displace this interaction whereas 10 mM glucose could not (Supplementary Fig. 2m). These data demonstrate that mItln2 binds mucus in a glycan-dependent manner.

We next asked if we could characterize the mucin binding profile of mItln2 using mucins isolated from tissues that support mItln2 expression. We tested mItln2 binding to MUC2, the predominant intestinal mucin in mammals; MUC5B, a salivary and bronchial mucin; and MUC5AC, a respiratory and gastrointestinal mucin that is induced during Th2 inflammation^35^. Purified porcine MUC2, MUC5AC, and MUC5B were immobilized to nitrocellulose, and assayed for mItln2 binding in the presence of calcium, EDTA, or 10 mM galactose. The lectin exhibited robust binding to all three mucins. These interactions were attenuated by the addition of EDTA or galactose competitor (Fig. 2e). These data suggest that mItln2 interacts with secreted mucins at its expression sites via recognition of Gal*p* residues.

Certain lectins, such as galectins and trefoil factors, crosslink mucin to strengthen the mucus barrier^36–38^. In examining mucin-mItln2 interactions by confocal microscopy, we observed that fluorescently labeled MUC5AC co-localized in large aggregates with fluorescently labeled mItln2, and that this aggregation was not observed in the absence of the lectin (Fig. 2f). These observations suggest that mItln2 crosslinks mucins. To examine this possibility directly, we utilized a kinetic spectroscopic assay to measure changes in light transmittance upon mixing lectin with mucin. Increasing amounts of mItln2 were titrated into solutions containing 0.01% MUC5AC, MUC5B, and MUC2 and evaluated for changes in percent transmission over time. We observed rapid and dose-dependent clustering of mucins by mItln2, which was abrogated by the addition of galactose (Fig. 2g; Supplementary Fig. 2n). Moreover, this activity was observed at concentrations similar to those previously reported for other mucus crosslinking lectins^36^. Using time-lapse confocal microscopy, we confirmed mucin crosslinking by mItln2; lectin–mucin complexes were observed to form and grow over time (Fig. 2h; Supplementary Fig. 2o). To test whether this crosslinking depends on carbohydrate binding, we used chemically defined mucin mimetic polymers displaying pendant β-lactose (a disaccharide with terminal β-Gal*p* residue)^39^. As with mucins, mItln2 promoted robust polymer clustering (Fig. 2i), further indicating that mItln2 binds and crosslinks mucins via carbohydrate recognition.

### MItln2 is a microbicidal lectin

The epitope β-Gal*p* is found widely across taxonomic classes of bacteria^40,41^. Therefore, we hypothesized that mItln2 would bind microbes. Flow-cytometry and microscopy experiments indicate that mItln2 robustly bound mouse fecal bacteria, confirming that it interacts with microbes in native communities (Fig. 3a,b)^13^. In line with this finding, mItln2 bound numerous gram-positive and gram-negative isolates from native mouse fecal communities, including *Limsolactobacillus reuteri, Lactobacillus murinus, Lactobacillus johnsonii, Bacillus paramycoides, Mucispirillum schaedleri* and pathobionts, including *Klebsiella pneumoniae*, methicillin-resistant *Staphylococcus aureus* (MRSA) and *Enterococcus faecalis* (Fig. 3c–e; Supplementary Fig. 3a–e). Moreover, mItln2 binding was calcium-dependent, an indication that it is glycan-mediated. Notably, mItln2 did not bind to all tested isolates, highlighting its specificity (Fig. 3f; Supplementary Fig. 3f–j). Together, these data demonstrate that mItln2 recognizes specific microbes that colonize tissues in which mItln2 is expressed.

**Figure 3:**
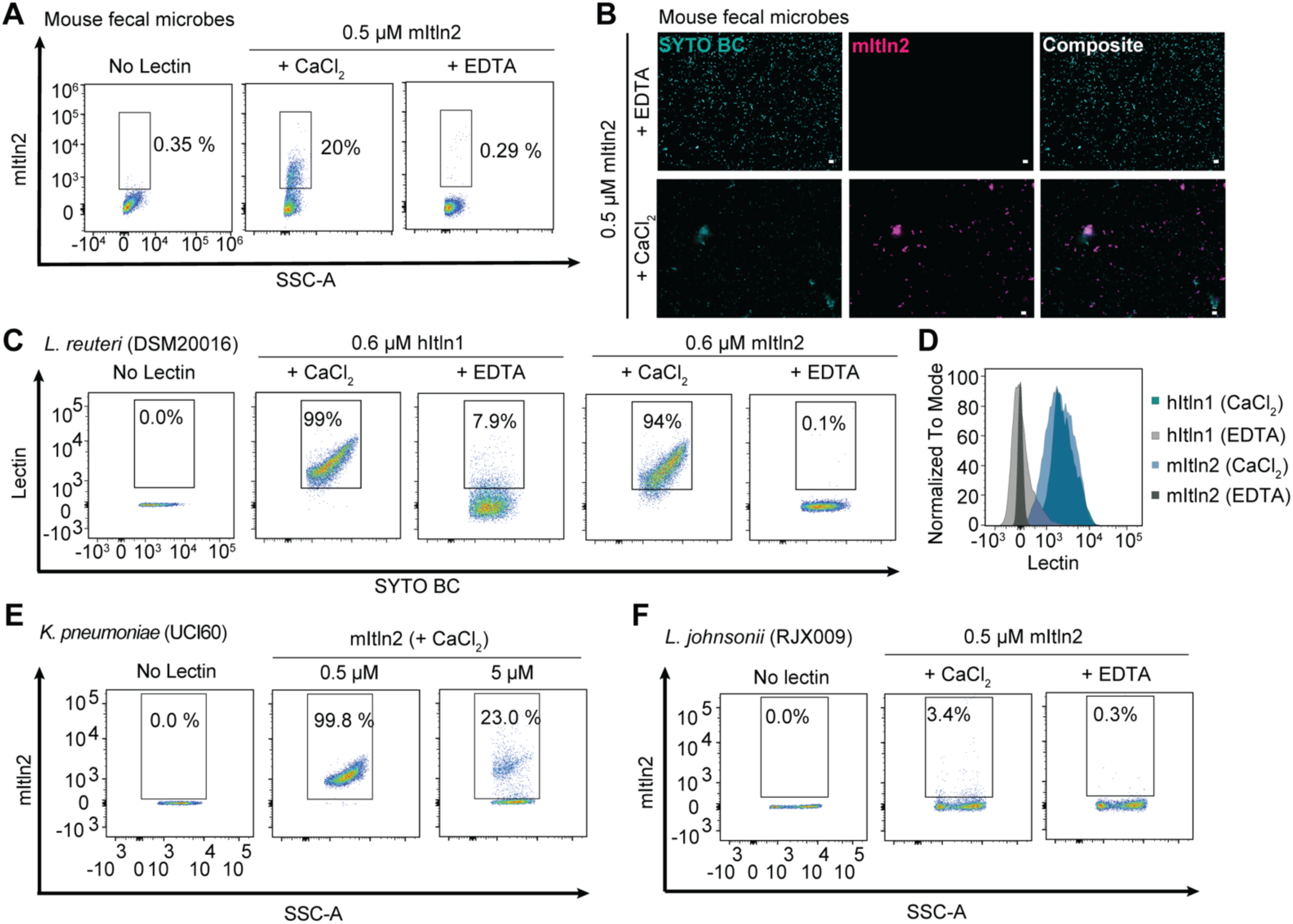
Determination of mItln2 binding to microbes. (**A**) Flow cytometry analysis of 0.5 μM mItln2 binding to mouse fecal samples under Ca^2+^ and EDTA conditions plotted as lectin binding (anti-Strep DY549) vs. SSC. (**B**) Images of murine fecal samples stained with 0.5 μM mItln2 (magenta) in the presence of EDTA (top) or Ca^2+^ (bottom) and counterstained with SYTO BC (teal). mItln2 was detected by Strep antibody. Scale bars, 10 μm. **(C** and **D**) Flow cytometry analysis of 0.6 μM hItln1 and mItln2 binding to *L. reuteri* under Ca^2+^ and EDTA conditions. Dot plots (C) show lectin binding (anti-Strep DY549) vs. nucleic acid stain (SYTO BC). Histogram (D) displays cell counts as a percent of the maximum signal against lectin binding in different conditions. (**E** and **F**) Flow cytometry analysis of mItln2 binding to *K. pneumoniae* UCI60 (E) and *L. johnonii* RJX009 (F) under Ca^2+^ or EDTA conditions. mItln2 was detected by Strep antibody. Unstained samples (no lectin) served as controls. Data in (A), (C), (D), and (E) are representative of three independent experiments. Data in (B) and (F) are representative of three independent experiments.

We next evaluated the consequences of mItln2 binding to microbes. Analysis of mItln2-bound microbes by flow cytometry revealed that mItln2 treatment caused marked shifts in forward scatter (FSC) versus side scatter (SSC) plot, in a region associated with debris (Fig. 4a,b; Supplementary Fig. 4a–e). This effect depended on the lectin concentration, with higher mItln2 concentrations resulting in increased events in the debris region for both gram-negative and gram-positive binders. This increase in debris was not observed in nonbinding isolates (Supplementary Fig. 4f–k) nor when the lectin binding was inhibited with EDTA (Supplementary Fig. 4b,c), suggesting that the effect is glycan-dependent. Moreover, treatment with equimolar amounts of hItln1 did not show this phenotype (Fig. 4a,b). Similar alterations in FSC vs. SSC profiles of microbes have been previously reported for other antimicrobial proteins, such as lysozyme, suggesting that mItln2 functions as an antimicrobial lectin^42^. Additionally, microscopy evaluation of mItln2-treated *L. reuteri* showed a loss of cell integrity and accumulation of ghost-like cells (empty bacterial cell envelopes) over time (Fig. 4c,d; Supplementary Movies 1 and 2). Together, these data suggest that mItln2 compromises the cellular integrity of bound bacteria and may influence cell viability.

**Figure 4:**
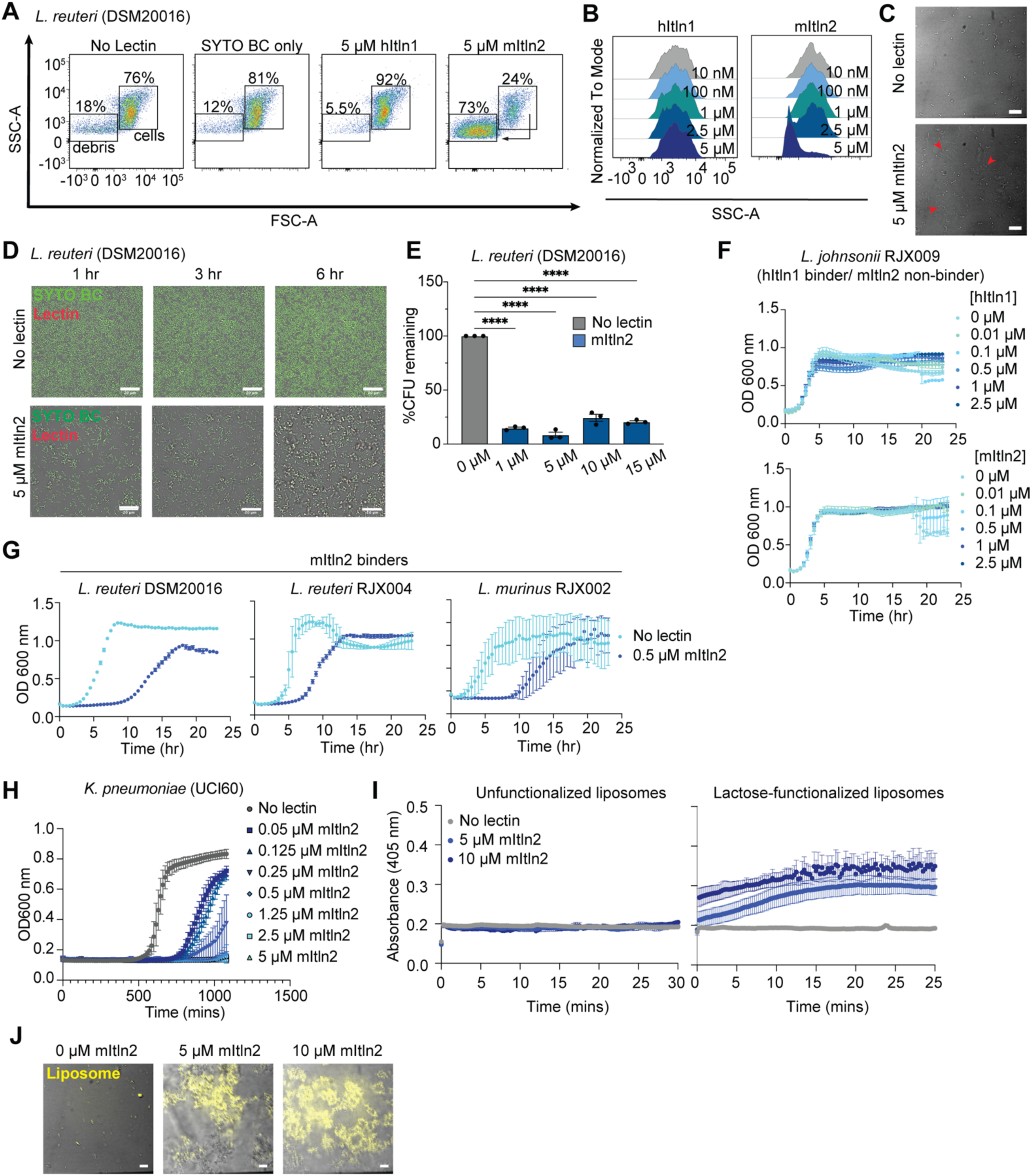
Assessment of mItln2’s impact on microbial viability. (**A**) Flow cytometry analysis of *L. reuteri* DSM20016 treated with 5 μM each of hItln1 and mItln2 in the presence of Ca^2+^ plotted as SSC-A vs. FSC-A. Regions representing debris and bacterial cells are indicated. Untreated bacteria and bacteria treated with SYTO BC served as controls. (**B**) Histogram plot displaying cell counts as a percent of the maximum signal against SSC-A for *L. reuteri* DSM20016 treated with various concentrations of hItln1 and mItln2. (**C**) Brightfield microscopy of *L. reuteri* DSM20016 after 3-hour treatment with 5 μM mItln2 in the presence of Ca^2+^. Examples of loss of cell integrity are indicated with red arrowheads. Untreated bacteria served as a control. Scale bars, 10 μm. (**D**) Time-lapse images of *L. reuteri* DSM20016 following treatment with 5 μM mItln2 (red) in the presence of Ca^2+^ and counterstained with SYTO BC (green). Scale bars, 20 μm. (**E**) Quantification of viable *L. reuteri* DSM20016 by dilution plating after incubation with various concentrations of mItln2 for 4 hours. Data show mean ± SEM (n = 3 independent experiments). ****P < 0.0001 (one-way ANOVA followed by Dunnett’s multiple comparisons test). (**F** and **G**) Recovery assay assessing microbial growth after lectin treatment. Microbes that bind hItln1 but not mItln2 (*L. johnsonii* RJX009) (F) and microbes that bind mItln2 (*L. reuteri* DSM20016, *L. reuteri* RJX004, and *L. murinus* RJX002) (G) were treated with the indicated lectins for 4 hours. Treated microbes were transferred to growth medium for recovery, with growth monitored by OD_600_ measurements. Untreated microbes served as controls. Data are shown as mean ± SD (n = 3 technical replicates). (**H**) Growth curve of *K. pneumoniae* UCI60 in the presence or absence of mItln2 at varying concentrations. Data show mean ± SEM (n=3 independent experiments). Untreated microbes served as controls. (**I** and **J**) Assessment of disruption of lactose-functionalized liposome, encapsulating texas red-dextran dye, after treatment with mItln2 for 1 hour. Spectroscopic assay (I) measured increase in absorbance at 405 nm following mItln2 treatment, with unfunctionalized liposomes serving as a control. Data are shown as mean ± SD (n = 3 technical replicates). Microscopy images (J) revealed dye leakage and loss of liposome integrity. Scale bars, 10 μm. Data in (C), (D), (F), (G), (I), and (J) are representative of two independent experiments. Data in (A) and (B) are representative of three independent experiments.

To assess whether mItln2 has microbicidal activity, we used a colony-forming unit (CFU) assay. Bacterial isolates were grown to mid-log phase and treated with increasing concentrations of mItln2. Consistent with the flow cytometry and microscopy data, a significant reduction in the viability of *L. reuteri* and *B. paramycoides* occurred upon treatment with low micromolar concentrations of mItln2 (Fig. 4e; Supplementary Fig. 4l). These concentrations are in line with other antimicrobial proteins shown to kill microbes^43,44^. The cell-killing effect of mItln2 was confirmed using a plate reader-based growth recovery assay, which showed a significant delay in the growth recovery for mItln2-binding *Lactobacillus* isolates following mItln2 treatment (Fig. 4f,g). Importantly, delayed growth recovery was not observed following treatment of unbound isolates with mItln2, nor in isolates recognized by hItln1 when treated with hItln1. These data reveal that mItln2 possesses microbicidal activity against specific gram-positive species, an effector function distinct from hItln1. To test the activity of mItln2 on gram-negative pathogenic bacteria, we used *K. pneumoniae*, a common opportunistic pathogen. We observed robust mItln2 binding and a shift in the FSC vs SSC plot for *K. pneumoniae* (Fig. 3e; Supplementary Fig. 4d). Furthermore, including mItln2 in the culture media led to dose-dependent growth inhibition of *K. pneumoniae*, which was observed even at low concentrations (50 nM) of mItln2 (Fig. 4h). These data suggest a mechanistic rationale for the finding that Th2 inflammation in BALB/c mice is protective against *K. pneumoniae* infection^45^. Collectively, the results indicate that mItln2 is a broad-spectrum antimicrobial protein that can target opportunistic pathogens at mucosal surfaces.

### MItln2 lyses bacterial cells

Other bactericidal mammalian lectins have been implicated in microbial membrane perturbation or pore formation, processes that can release intracellular contents^43,44^. Given that mItln2 alters bacterial morphology and reduces viability, we postulated that mItln2 acts as a cytolytic agent by disrupting microbial membranes. In support of this hypothesis, we observed loss of DNA signal in bacteria stained with DNA intercalating dye (SYTO BC) following mItln2 treatment (Supplementary Fig. 5a; Supplementary Movie 3). We also found an increased uptake of propidium iodide, a DNA dye that cannot penetrate intact membranes (Supplementary Fig. 5b).

We employed a liposome disruption assay to better understand how mItln2 exerts its microbicidal activity (Supplementary Fig. 5c). In this approach, we generated 200 nm diameter liposomes composed of palmitoyl oleoyl phosphatidylcholine (POPC), 10 mol% cholesterol, and 10 mol% 1,2-dioleoyl-sn-glycero-3-phosphethanolamine-N-dibenzocyclooctyl (18:1 DBCO PE) as a functional handle to append specific glycans. Liposomes were functionalized with β-lactose (a disaccharide with terminal β-Gal*p* residue) and treated with increasing concentrations of lectins^46^. Treatment of lactose-functionalized liposomes with mItln2 led to a concentration-dependent increase in absorbance at 405 nm, indicative of membrane disruption (Fig. 4i). This process was glycan-dependent, as adding mItln2 to liposomes lacking a glycan ligand afforded no change in absorbance. Confocal microscopy further confirmed the glycan-dependent ability of mItln2 to disintegrate the lactose-functionalized liposomes encapsulating Texas red-dextran dye (Fig. 4j). Moreover, although mItln2 bound mammalian cells, propidium iodide uptake assays demonstrate that it did not compromise cellular membrane integrity, indicating that mItln2 specifically diminishes microbial viability (Supplementary Fig. 5d,e).

Unlike other cytolytic proteins reported, the disruption of liposomes by mItln2 depended on glycan binding and occurred at physiological pH (7.4) and osmolarity (150 mM NaCl). The liposome disruption and the cell-killing activity did not require acidic pH or low salt concentration (<50 mM)^43,47–49^. These findings reveal that the powerful cell-killing ability of mItln2 synergizes with its ability to bind and cluster mucins. Thus, mItln2 has dual roles during Th2 inflammation: it functions by crosslinking host mucins and by binding and killing microbes.

### Human intelectin-2 (hItln2) is a galactose-binding antimicrobial lectin

Human intelectin-2 (hItln2) is constitutively expressed by Paneth cells in the small intestine and is detectable in the lungs^19,50^. A recent study demonstrated that hItln2 expression is even higher than hItln1 in healthy small intestinal tissue and that aberrant expression of hItln2 is observed in both the ileum and colon of CD patients^19^.Consistent with its constitutive intestinal secretion, hItln2 is readily detected in human fecal samples (Supplementary Fig. 6a). However, as with mItln2, the ligand specificity and biological function of hItln2 have not been clear.

To characterize hItln2, we expressed and purified recombinant StrepII-tagged hItln2. The production of recombinant hItln2 required optimization, particularly through the introduction of a glycosylation site (A177S) and the removal of an unpaired cysteine near the C-terminus (C311G) (Supplementary Fig. 6b). These modifications improved solubility and reduced aggregation, yielding soluble, stably folded protein with a melting temperature of approximately 54 °C (Supplementary Fig. 6C). We also observed that hItln2 formed covalent trimers and higher-order oligomers, similar to native hItln2 in small intestinal tissue (Supplementary Fig. 6d,e)^19^.

Like mItln2, the ligand binding site of hItln2 lacks the aromatic box present in Itln1 (Fig. 5A). In hItln2, a serine at position 309 replaces the tyrosine at the corresponding site in hItln1. We reasoned that this more open binding pocket might accommodate pyranose sugars, as observed with mItln2. We next investigated if hItln2 and mItln2 share glycan targets. Using BLI to evaluate the binding of hItln2 with immobilized mono- and di- saccharides, we observed that hItln2 indeed binds mItln2 ligands (β-Gal*p* and β-6SO_3_ LacNAc) but not the Itln1 ligand (β-Gal*f*) (Fig. 5b–d). The apparent dissociation constant of hItln2 for β-Gal*p* was around 3.9 μM (Supplementary Fig. 6f,g). Thus, mItln2 and hItln2 share common ligands. Moreover, the microbial glycan array analysis for hItln2 yielded hits containing Gal*p*, but also phosphoethanolamine (PEt), suggesting that PEt may serve as an additional microbial ligand for hItln2 (Fig. 5e,f; Supplementary Table 3). These overlapping yet distinct ligand recognition properties could be due to other differences in the binding site residues of mItln2 and hItln2 (mItln2/hItln2: E274/Q286, A288/W300, and Y297/S309).

**Figure 5:**
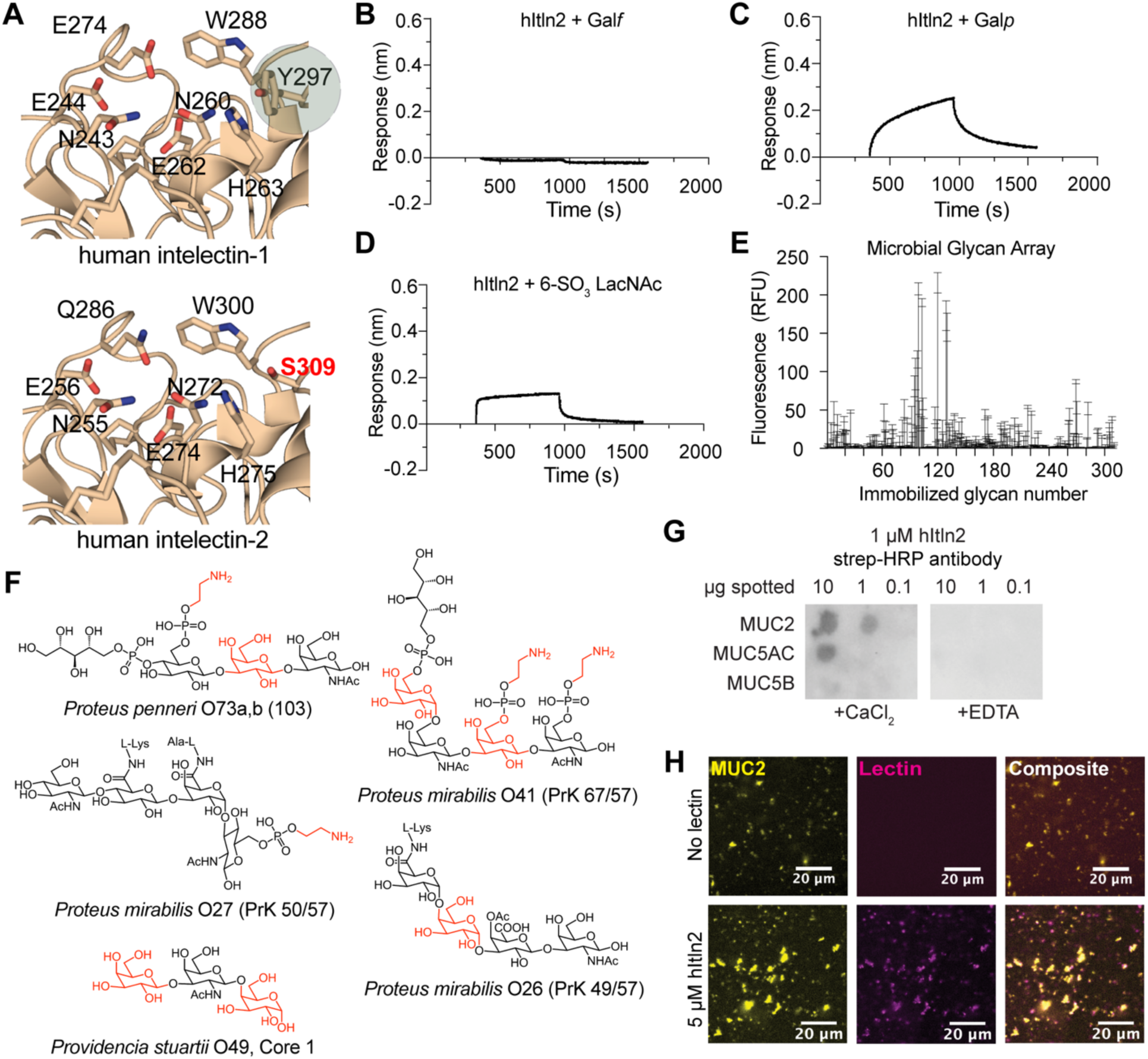
Characterization of glycan specificity for hItln2. (**A**) Comparison of carbohydrate recognition domain of hItln1 (top, PDB ID 4WMY) and hItln2 (bottom, predicted model), showing conserved calcium coordination residues and a serine replacing tyrosine at position 309 in hItln2. (**B** – **D**) BLI traces of hItln2 binding to immobilized biotinylated-β-Gal*f* (a hItln1 ligand), (B), biotinylated-β-Gal*p* (C), and biotinylated-6SO_3_ LacNAc (D). In (B) and (C), 1.5 μM protein was used. In (D), 10 μM protein was used. Data were normalized by background subtraction, using biotin-loaded streptavidin. (**E**) Binding of recombinant hItln2 (25 µg/mL) to a microbial glycan microarray from NCFG. Data are shown as mean ± SD (n = 4 technical replicates). (**F**) Glycan structures bound by hItln2 in glycan arrays. All hits contain either a Gal*p* or ethanolamine, shown in red. (**G**) Dot blot analysis of MUC2, MUC5AC, and MUC5B, spotted on the membrane and probed with 1 μM hItln2 in the presence of Ca^2+^ or EDTA. Binding of hItln2 was detected using a Strep-HRP antibody. (**H**) Images of 5 μM hItln2 (magenta) binding to 0.01% (w/v) fluorescently labeled MUC2 (yellow). Binding of hItln2 was detected with Strep antibody. Scale bars, 20 µm. Results in (B), (C), (D) and (G) are representative of three independent experiments. Result in (H) is representative of two independent experiments.

Because mItln2 and hItln2 both bind to β-Gal*p*, we reasoned that they might interact with similar physiological substrates. Supporting this hypothesis, we found that hItln2 binds intestinal mucins, MUC2, and MUC5AC in a calcium-dependent manner (Fig. 5g,h). Spectroscopic analysis of hItln2–mucin interactions showed mucin crosslinking by hItln2, but hItln2 activity was more challenging to detect via microscopy (Supplementary Fig. 6h–j). Additionally, hItln2 recognized microbes within a human fecal sample (Fig. 6a,b), as well as a range of gram-positive and gram-negative bacterial isolates that includes the opportunistic pathogens *Escherichia coli* and *Enterococcus faecalis*. These cell-binding properties were inhibited by the chelator EDTA (Supplementary Fig. 7a–f), suggesting that hItln2 interacts with glycans through its calcium ion. Unlike mItln2, hItln2-treated microbes formed large cellular aggregates, which could be disrupted with EDTA, suggesting glycan-dependent agglutination (Supplementary Fig. 7b and d).

**Figure 6:**
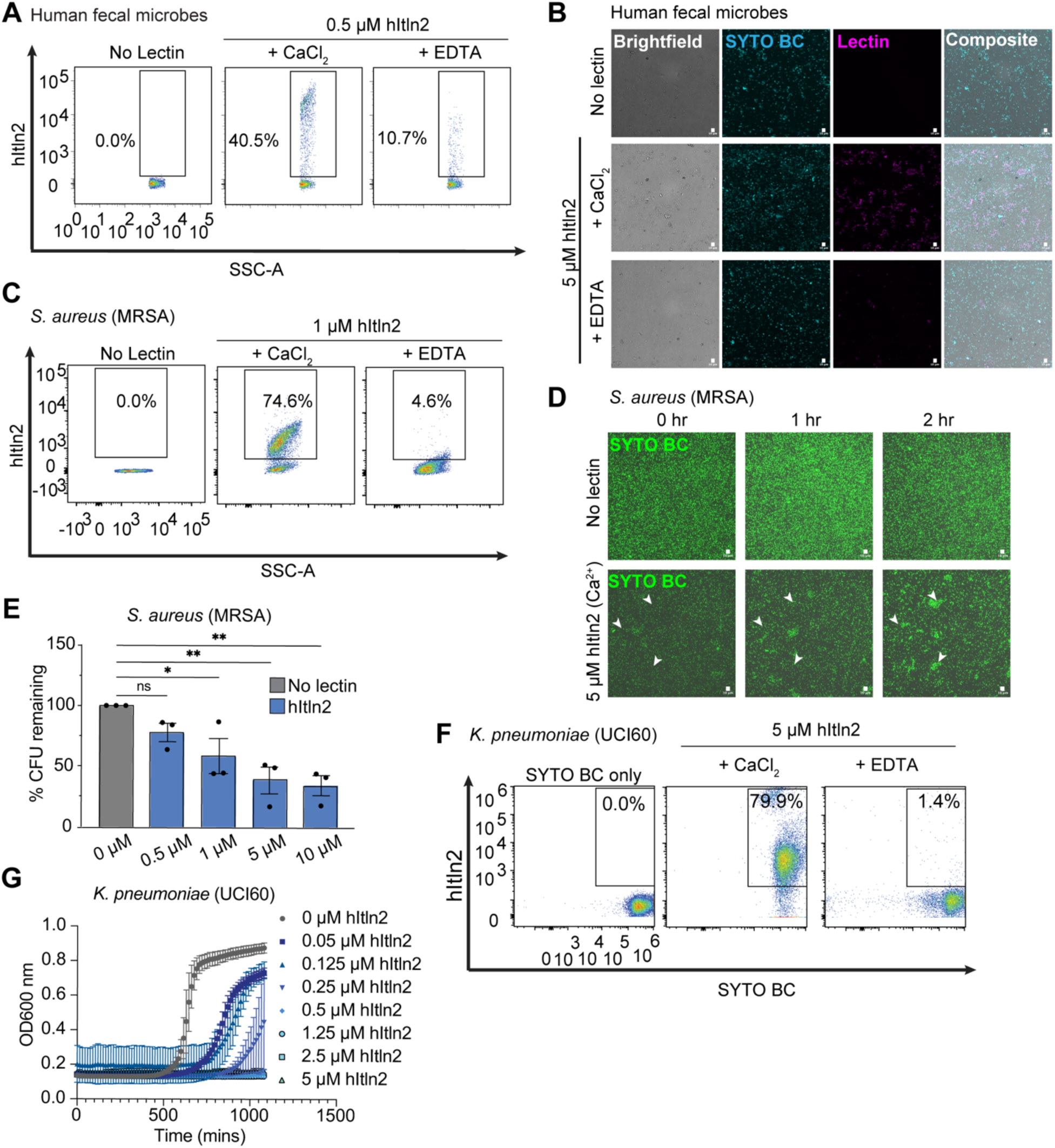
Evaluation of hItln2 binding to microbes and its impact on microbial viability. (**A**) Flow cytometry analysis of 0.5 μM hItln2 binding to human fecal samples under Ca^2+^ and EDTA conditions, plotted as lectin binding (Streptactin DY549) vs. SSC**). (B)** Microscopy images of human fecal samples stained with 5 μM hItln2 (magenta) in the presence of Ca^2+^ or EDTA and counterstained with SYTO BC (teal). hItln2 was detected by Streptactin. Scale bars, 10 µm. (**C**) Flow cytometry analysis of 1 μM hItln2 binding to *S. aureus* MRSA under Ca^2+^ and EDTA conditions, plotted as lectin binding (anti-Strep DY549) vs. SSC. Treatment with SYTO BC and Strep antibody (without lectin) served as a control. (**D**) Time-lapse images of *S. aureus* MRSA treated with 5 μM hItln2 in the presence of Ca^2+^ and counterstained with SYTO BC (green). Untreated bacteria served as a control. Examples of cell agglutination are indicated with white arrowheads. Untreated bacteria served as control. Scale bars, 10 µm. (**E**) Viable *S. aureus* MRSA quantified by dilution plating after incubation with various concentrations of hItln2 for 4 hours. Data represent mean ± SEM (n = 3 independent experiments). ns: not significant, *P < 0.05, **P < 0.01 (one-way ANOVA followed by Dunnett’s multiple comparisons test). (**F**) Flow cytometry analysis of hItln2 binding to *K. pneumoniae* UCI60 under Ca^2+^ or EDTA conditions, detected by Streptactin. Treatment with SYTO BC (without lectin) served as a control. (**G**) Growth curve of *K. pneumoniae* UCI60 in the presence or absence of hItln2 at varying concentrations. Data show mean ± SEM (n=3 independent experiments). Data in (A), (B), (C), (D), and (F) are representative of three independent experiments.

We next sought to investigate if hItln2 functions as an antimicrobial protein based on several key characteristics: 1) its abundant expression at sites of microbial colonization; 2) its secretion from canonically anti-microbial Paneth cells; and 3) its cationic nature (isoelectric point of 8.3) and 4) its lengthened N-terminal sequence (relative to hItln1) that is predicted to form an alpha-helix (Supplementary Fig. 6k). We first evaluated if hItln2 could bind the pathobiont *Staphylococcus aureus*, a common Gram-positive opportunistic pathogen. Increasing evidence suggests that *S. aureus* can colonize the human gut and, under the right conditions, can disseminate to other host tissues via translocation across the intestinal mucosa and epithelium^51^. We reasoned that hItln2 might encounter this organism in the gastrointestinal tract. Consistent with this hypothesis, hItln2 showed robust binding to and pronounced agglutination of methicillin-resistant S. aureus (MRSA) by flow cytometry and microscopy (Fig. 5c,d).

To evaluate the antimicrobial properties of hItln2, we performed CFU assays on hItln2-treated *S. aureus*. For this, MRSA was grown to mid-log phase and then treated with increasing amounts of hItln2. Following treatment, cells were vigorously mixed, serially diluted in assay buffer with thorough mixing between dilutions, and plated on growth substrate, and allowed to grow overnight at 37 °C before counting colonies. Similar to mItln2, hItln2 treatment decreased CFU counts for MRSA, indicating reduced bacterial viability (Fig. 6e). This result was also observed after treatment of other gram-positive bacteria recognized by hItln2 (Supplementary Fig. 7h). HItln2 also bound and inhibited the growth of the gram-negative pathobiont *K. pneumoniae* in a dose-dependent manner (Fig. 6f,g). Together, these data suggest that hItln2 inhibits microbial viability and growth through the antimicrobial effector function of agglutination. Like mItln2, hItln2 did not compromise the membrane integrity of mammalian cells (Supplementary Fig. 7i,j).

Expression of hItln2 in healthy subjects is primarily restricted to the small intestine^19^. Our initial characterization of hItln2 activity was performed under ileum-like conditions: neutral pH and physiological ionic strength. However, hItln2 is also produced in the colons of patients with colonic CD and UC due to Paneth cell metaplasia^19^. The colonic environment in CD and UC patients can range from acidic to neutral pH (pH 4-7) and has altered ionic strength owing to aberrations in electrolyte uptake and secretion during inflammation^52–54^. Moreover, other small intestinal-derived antimicrobial proteins are reported to be active only under acidic pH and low ionic strength^43,47–49^. To address these variables, we next investigated the activity of hItln2 at low pH and ionic strength.

Even at low pH and ionic strength, hItln2 remained well-folded and retained binding to β-Gal*p* and β-6SO_3_ LacNAc (Fig. 7a; Supplementary Fig. 8a,b). We also confirmed that hItln2 bound to mucins and microbes under these conditions (Fig.7b–d; Supplementary Fig. 8c–e); however, EDTA addition did not abrogate these interactions, suggesting a shift to an alternate binding mode. Moreover, under these conditions, hItln2 mediated rapid and robust formation of mucin and microbial aggregates (Fig. 7e,f; Supplementary Fig. 8f,g). Similarly, CFU analysis showed a dramatic reduction in the growth of the pathobionts, *S. aureus*, *B. paramycoides*, and *E. coli*, confirming that the agglutination-mediated antimicrobial function of hItln2 was enhanced under conditions that represent intestinal inflammation (Fig. 7g,h; Supplementary Fig. 7h–j).

**Figure 7:**
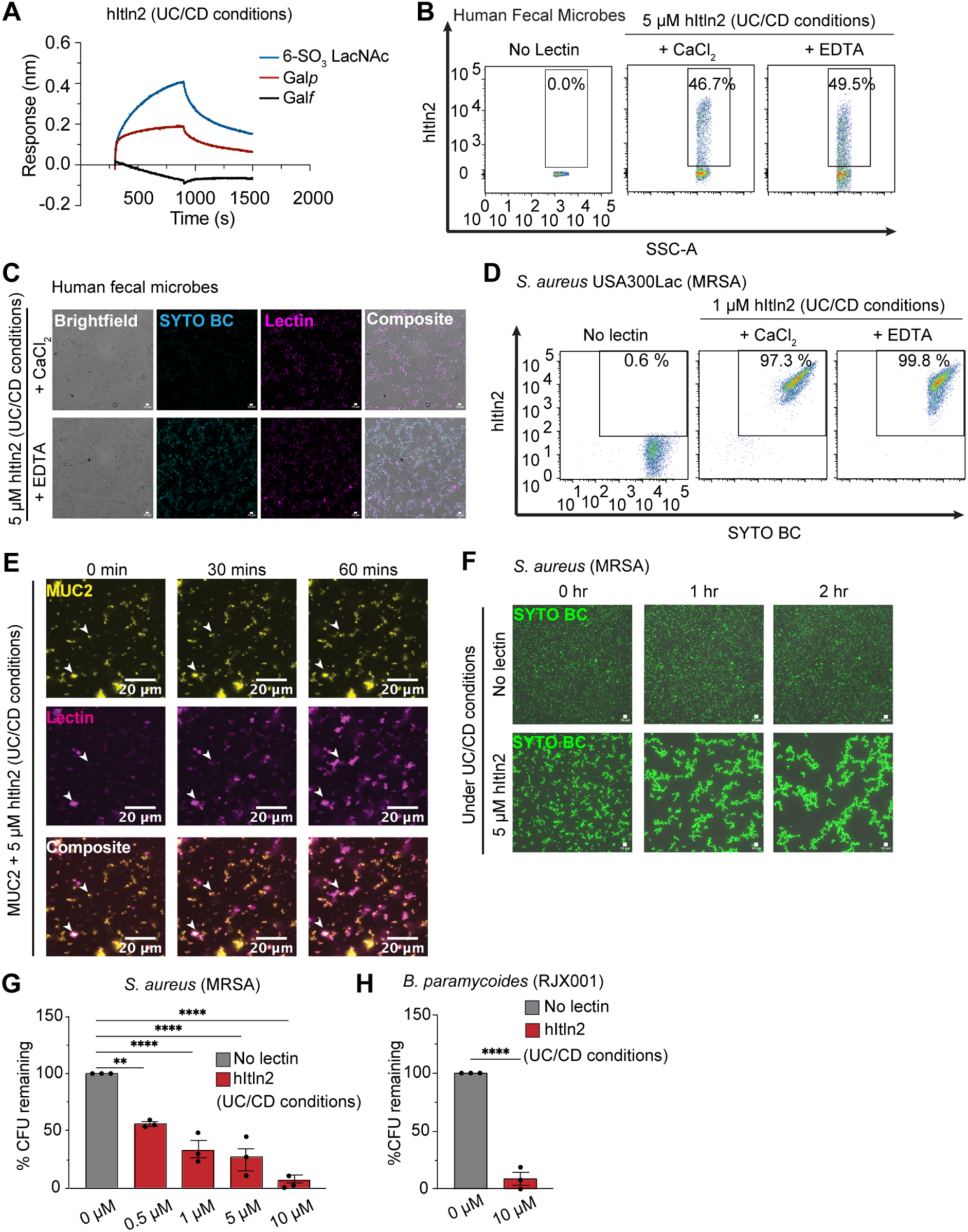
Evaluation of hItln2 binding to mucin and microbes at low pH and low salt (UC/CD conditions) (**A**) BLI trace of 1.5 μM hItln2 binding to immobilized biotinylated-β-Gal*f*, biotinylated-β-Gal*p,* and biotinylated-6SO_3_ LacNAc under low pH and salt conditions that mimic the intestinal environment during active UC or CD. Data were normalized by background subtraction, using biotin-loaded streptavidin. (**B** and **C**) Flow cytometry (B) and microscopy (C) of 5 μM hItln2 (UC/CD) binding to human fecal samples in Ca^2+^ and EDTA conditions under low pH and salt concentration (UC/CD conditions). Dot plot (B) displays lectin binding (Streptactin DY549) vs. SSC. hItln2 was detected by Streptactin. (**D)** Flow cytometry of 1 μM hItln2 (UC/CD) binding to *S. aureus* MRSA in Ca^2+^ and EDTA conditions under low pH and salt concentrations (UC/CD conditions), detected by Streptactin. Treatment with SYTO BC and Streptactin (without lectin) served as a control. (**E**) Time-lapse images of 0.01% (w/v) fluorescently labeled MUC2 (yellow) treated with 5 μM hItln2 (magenta) under low pH and low salt (UC/CD) conditions. Binding of hItln2 was detected with Strep antibody. Examples of mucin crosslinking are indicated with white arrowheads. Scale bars, 20 µm. (**F**) Time-lapse images of *S. aureus* MRSA treated with 5 μM hItln2 in the presence of Ca^2+^ and counterstained with SYTO BC (green) under low pH and low salt (UC/CD) conditions. Untreated bacteria served as a control. Scale bars, 10 µm. (**G**) Viable *S. aureus* MRSA quantified by dilution plating after incubation with various concentrations of hItln2 for 4 hours under low pH and low salt (UC/CD) conditions. Data show mean ± SEM (n = 3 independent experiments). **P < 0.01, ****P < 0.0001 (one-way ANOVA followed by Dunnett’s multiple comparisons test). (**H**) Quantification of viable *B. paramycoides* RJX001 by dilution plating after treatment with 10 μM hItln2 for 4 hours at low pH and low salt (UC/CD) conditions. Data represent men ± SEM (n = 3 independent experiments). ****P < 0.0001 (unpaired two-tailed t-test). Data in (A) and (E) are representative of two independent experiments. Data in (B), (C), (D), and (F) are representative of three independent experiments.

## DISCUSSION

Both mItln2 and hItln2 have been linked to infection and disease^6,7,19,33^; however, their specific effector functions were unclear. Here, we identified these proteins as galactose-binding lectins with protective functions in host defense against microbes. Specifically, our findings reveal that mItln2 and hItln2 recognize Gal*p*-containing glycans, conferring two key functional properties: mucin crosslinking and antimicrobial activity.

The sequence similarity between Itln1 and Itln2 indicates that they share similar overall structures. Itln1 binds exocyclic vicinal diols on microbial carbohydrates, like Gal*f*, through a calcium ion, with aromatic residues surrounding the saccharide. However, Itln1 does not bind the Itln2 ligand Gal*p*, which lacks an exocyclic vicinal diol. A critical difference between the Itln1 and predicted Itln2 binding sites is the absence of an aromatic residue in Itln2. Substitution at this position removes the aromatic box and results in a more open binding site, which could facilitate binding of Itln2 to vicinal diols within the pyranose ring of galactose. These findings highlight the versatility of the intelectin fold, where changing a single residue in the binding site alters glycan specificity. This feature suggests intelectins are excellent platforms for engineering lectins with unique binding specificities.

In mice, Th2 cytokines elicit the conditional expression of mItln2, suggesting its inducibility in response to pathogens capable of breaching mucus barriers. The ability of mItln2 to crosslink mucin and kill microbes highlights its protective role in strengthening the mucosal barrier and limiting the spread of infection during Th2 immunity. The ability of mItln2 to kill both pathogens and commensals may explain its limited and inducible expression in healthy tissues, where commensals are essential for maintaining host health. In contrast to some antimicrobial lectins that only exert antimicrobial activity under low pH and low salt conditions and in a glycan-independent manner, the bactericidal effect of mItln2 is distinctive due to its glycan dependence and efficacy under physiological pH and osmolarity conditions^43^. Moreover, we found that mItln2 did not affect mammalian cell viability, which suggests a possible role of membrane composition on lectin activity. These characteristics underscore the selectivity and robust antimicrobial function of mItln2, which is necessary for its activity at sites beyond the gastrointestinal tract, such as the lungs.

Although it has been argued that hItln2 is not an ortholog of mItln2^19^, these homologous proteins share the ability to crosslink host mucins. We found that the antimicrobial function of hItln2 involves microbial agglutination and inhibition of bacterial growth. Microbial agglutination was not observed with mItln2 but is a functional activity of other antimicrobial proteins, including zymogen granule proteins and human α-defensin-6^55–57^. Under homeostatic conditions in the small intestine, constitutively expressed hItln2 might serve to prevent the overgrowth of microbes via agglutination. However, in environments with higher acidity and altered ionic strength, as observed in CD and UC patients, ligand recognition by hItln2 occurs independent of calcium, and its mucin-crosslinking and antimicrobial activities are significantly enhanced. This change in activity suggests that hItln2 may act as a sensory switch. Moreover, these findings highlight how the lectin function can be altered by inflammation. Still, it remains unclear whether the enhanced activities of hItln2 under pathological conditions help limit inflammation and reestablish mucosal health or if they alter the microbial composition and exacerbate inflammation. Further investigations are needed to distinguish between these roles.

Our data support a model in which Itln2 protects the host against microbial threats: a defensive role in which it binds mucins to reinforce the mucosal environment and an offensive role in which it reduces bacterial burden via its antimicrobial activities. These findings advance our understanding of intelectins as crucial players in host–microbe interactions, with implications for maintaining mucosal health in both homeostatic and diseased states.

## ACKNOWLEDGMENTS

The authors thank the MIT Biophysical Instrumentation Facility, Drs. Bradley Turner and Mike Wuo, for assistance with circular dichroism and biolayer interferometry. We thank Drs. Robert Kerby and Federico Rey for *L. reuteri* DSM20016 strain, Dr. Edward Hollox for discussions on the B6.C-Itln1-6 mouse model, and Prof. Eric Alm for generously providing human fecal samples. The authors thank the Koch Institute’s Robert A. Swanson (1969) Biotechnology Center for microscopy and flow cytometry core facilities. The research reported in this publication was supported by NIH Glycoscience Common Fund U01 CA231079 (L.L.K), NIAID R01 AI055258 (L.L.K), U01 AI125926 (C.L.B), R37 AI32738 (C.L.B.), NIDDK P30 DK043351 (R.J.X.), NSF EF-2125118 (K.R.), and ICB-2024-BEM-10 (under award W911NF1920026) (K.R.). A.E.D. thanks Dr. Kittikhun Wangkanont for helpful discussions.

## AUTHOR CONTRIBUTIONS

Conceptualization: AED, DS, EBN, CLB, and LLK; Methodology: AED, DS, EBN, RSC, ALP, DS, CD, JWA, CEB, MK, and GCO; Investigation: AED, DS, EBN, RSC, ALP, JJY, JI, DS, CD, SB, SJ, CE, SG, JWA, CEB, MK, and GCO; Visualization: AED, DS, EBN, RSC, ALP; Funding acquisition: KR, RJX, CLB, and LLK; Project administration: GP, HV, KR, RJX, CLB, and LLK; Supervision: KR, RJX, CLB, and LLK; Writing – original draft: AED, DS, EBN, CLB, and LLK; Writing – review & editing: AED, DS., EBN, RSC, ALP, SG, RX, CLB, and LLK

## CORRESPONDANCE

Further information and requests for resources and reagents should be directed to and will be fulfilled by the lead contact, Laura L. Kiessling.

