## Supporting Information for "Intelectin-2 is a broad-spectrum antimicrobial lectin"

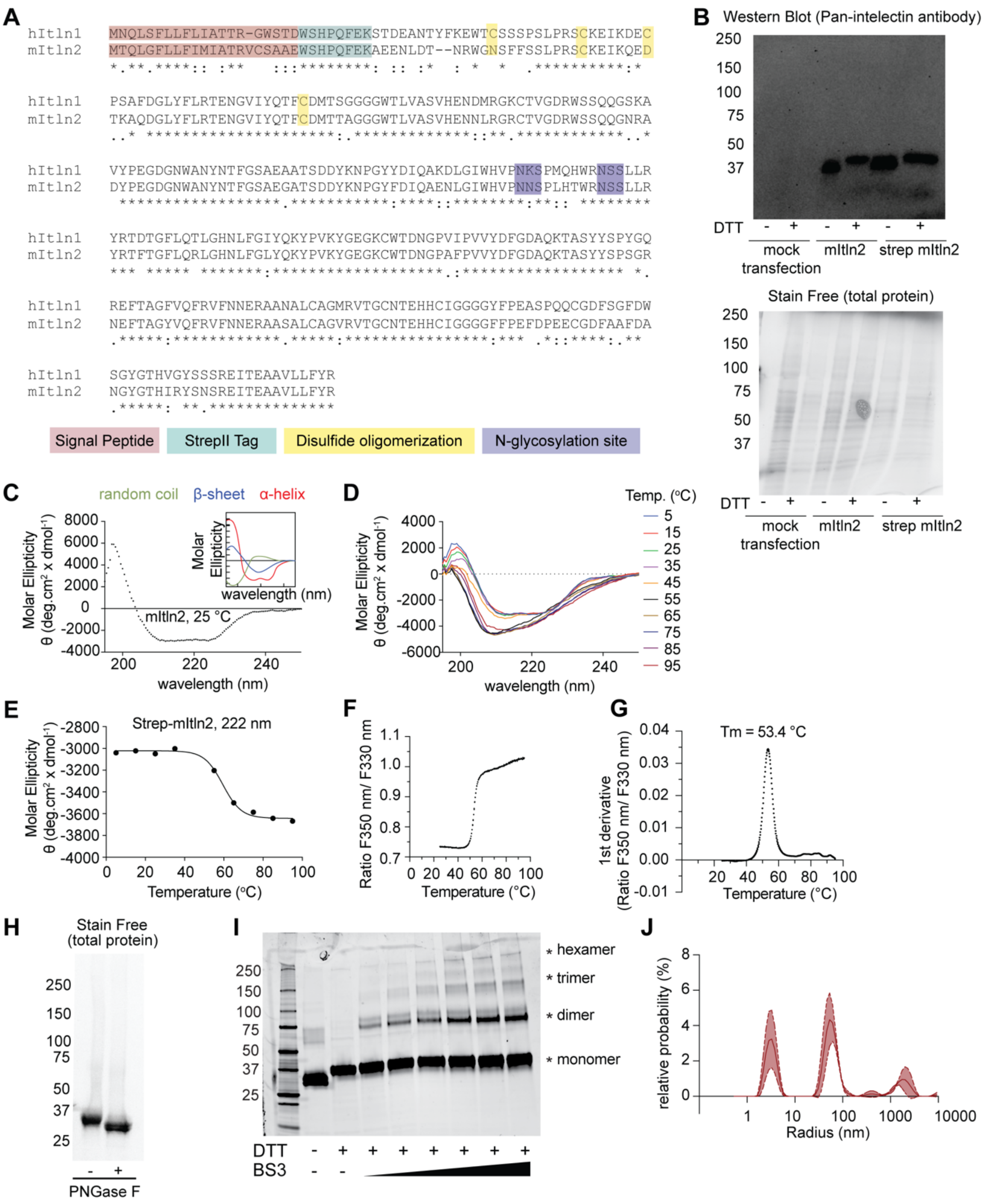
A
hItln1 MNQLSFLLFLIATTR-GWSTDWSHPQFEKSTDEANTYFKEWTCSSSPSLPRSCKEIKDEC
mItln2 MTQLGFLLFIMIATRVCSAAEWSHPQFEKAEENLDT--NRWGNSFFSSLPRSCKEIKQED
Signal Peptide
StrepII Tag
Disulfide oligomerization
N-glycosylation site
B
Western Blot (Pan-intelectin antibody)
Stain Free (total protein)
DTT
mock transfection
mItln2
strep mItln2
C
random coil
β-sheet
α-helix
Molar Ellipticity θ (deg.cm² x dmol⁻¹)
mItln2, 25 °C
wavelength (nm)
D
Temp. (°C)
E
Strep-mItln2, 222 nm
Temperature (°C)
F
Ratio F350 nm/ F330 nm
G
Tm = 53.4 °C
1st derivative (Ratio F350 nm/ F330 nm)
H
Stain Free (total protein)
PNGase F
I
hexamer
trimer
dimer
monomer
DTT
BS3
J
relative probability (%)
Radius (nm)

**Supplementary Figure 1: Expression and physical characterization of recombinant mItln2.**

(**A**) MUSCLE sequence alignment of StrepII-tagged hItln1 and StrepII-tagged mItln2. Signal peptide (pink), StrepII-tag (cyan), and N-glycosylation motifs (purple) are highlighted. Cysteine residues needed for oligomerization of hItln1 but absent in mItln2 are indicated (yellow). Conserved, highly similar, and similar residues are represented by "*", ":", and "." respectively. (**B**) Western blot (top) and corresponding stain-free gel (bottom) of the culture medium from HEK293T cells transfected with native mItln2 (no-tag) or StrepII-tagged mItln2 run under reducing and non-reducing conditions. Recombinant mItln2 was detected using a pan-intelectin antibody. (**C** and **D**) Circular dichroism spectra of mItln2 protein (500 μg/mL in PBS buffer) from 180 nm to 260 nm at different temperatures. Standard spectra of the basic secondary structures of a polypeptide chain (α-helix, β-sheet, and random coil) are embedded in (C). (**E**) Changes in ellipticity of mItln2 protein at 222 nm as a function of temperature, derived from data in (D). (**F** and **G**) Ratio of change in intrinsic fluorescence intensity at 330 nm and 350 nm (F) and its first derivative (G) as a function of temperature for recombinant mItln2, obtained from differential scanning fluorimetry. F, fluorescence. (**H**) Stain-free gel of untreated and PNGase F-treated mItln2 proteins under reducing condition, demonstrating a decrease in molecular weight after removal of glycans. (**I**) SYPRO Ruby-stained SDS-PAGE of mItln2 proteins (1 mg/ mL) crosslinked with various concentrations of bis(sulfosuccinimidyl)suberate (BS3: 0.1 mM, 0.25 mM, 0.5 mM, 1.25 mM, 2.5 mM, and 5 mM). Samples were run under either reducing or non-reducing conditions as indicated. Molecular weight of monomeric mItln2 is 34 kDa, and different oligomers are indicated by *. (**J**) Relative distributions of particle sizes for mItln2 protein, obtained from dynamic light scattering. Results shown in (B), (C), (D), (E) are representatives of two independent experiments. Results shown in (F), (G), (H), (I), and (J) are representatives of three independent experiments.

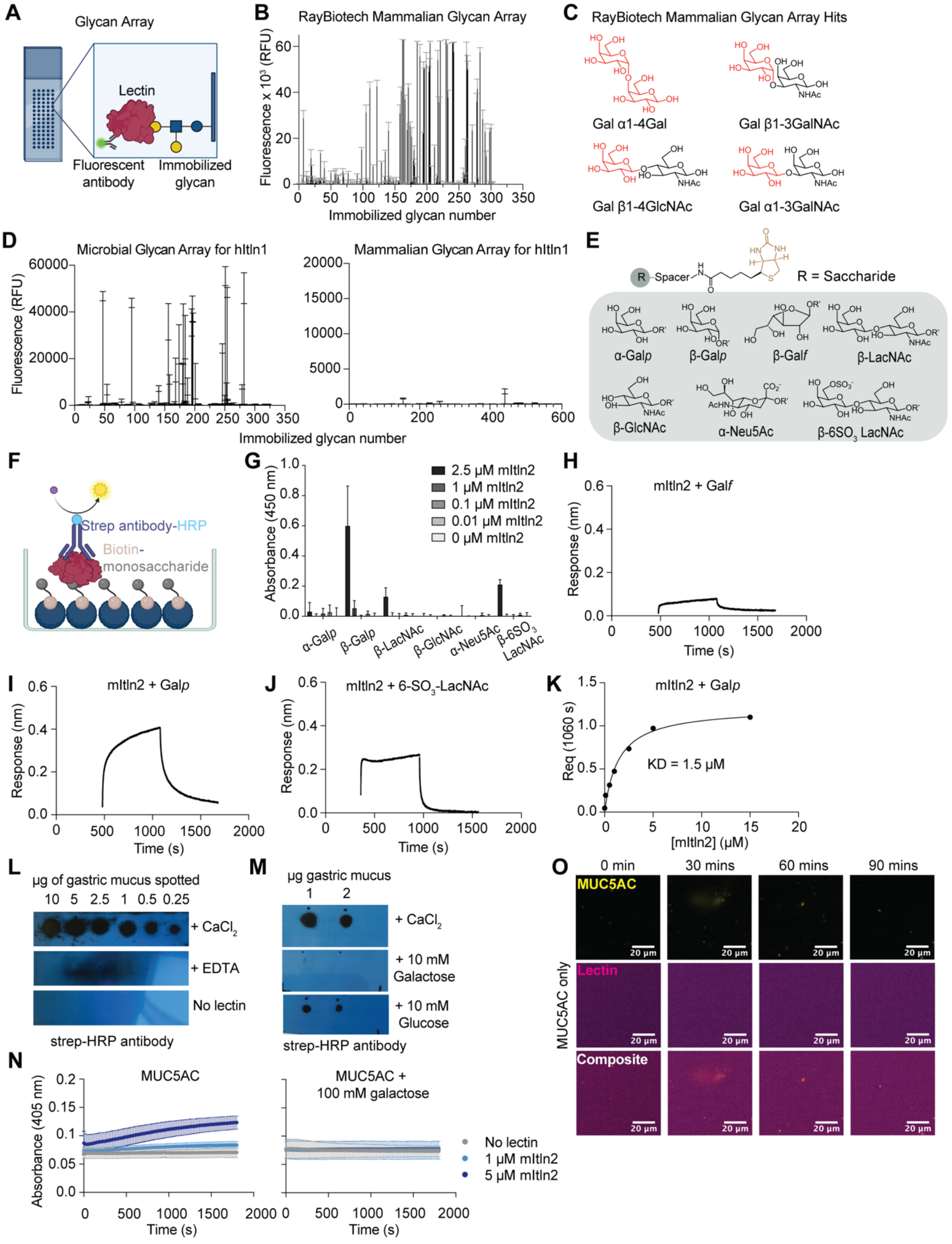
A
Glycan Array
Lectin
Fluorescent antibody
Immobilized glycan
B
RayBiotech Mammalian Glycan Array
Fluorescence x 10³ (RFU)
Immobilized glycan number
C
RayBiotech Mammalian Glycan Array Hits
Gal α1-4Gal
Gal β1-3GalNAc
Gal β1-4GlcNAc
Gal α1-3GalNAc
D
Microbial Glycan Array for hItln1
Mammalian Glycan Array for hItln1
Fluorescence (RFU)
Immobilized glycan number
E
R = Saccharide
α-Galp
β-Galp
β-Galf
β-LacNAc
β-GlcNAc
α-Neu5Ac
β-6SO3 LacNAc
F
Strep antibody-HRP
Biotin-monosaccharide
G
Absorbance (450 nm)
2.5 μM mItln2
1 μM mItln2
0.1 μM mItln2
0.01 μM mItln2
0 μM mItln2
H
mItln2 + Galf
Response (nm)
Time (s)
I
mItln2 + Galp
J
mItln2 + 6-SO3-LacNAc
K
mItln2 + Galp
Req (1060 s)
KD = 1.5 μM
[mItln2] (μM)
L
μg of gastric mucus spotted
10 5 2.5 1 0.5 0.25
+ CaCl2
+ EDTA
No lectin
strep-HRP antibody
M
μg gastric mucus
1 2
+ CaCl2
+ 10 mM Galactose
+ 10 mM Glucose
strep-HRP antibody
N
MUC5AC
MUC5AC + 100 mM galactose
Absorbance (405 nm)
Time (s)
No lectin
1 μM mItln2
5 μM mItln2
O
0 min
30 mins
60 mins
90 mins
MUC5AC
Lectin
Composite
MUC5AC only
20 μm

**Supplementary Fig 2: Carbohydrate binding specificity of recombinant mItln2.**
(A) Schematic of glycan array analysis. Recombinant lectins are allowed to bind synthetic and natural glycans immobilized on a glass slide. Lectin binding is detected with a fluorescent antibody against the lectin. (B) Binding profile of recombinant mItln2 (25 μg/mL) to mammalian glycan microarrays from RayBiotech. Data are shown as mean ± SD (n = 4 technical replicates). (C) Glycan structures recognized by mItln2 in mammalian glycan microarrays from RayBiotech. All hits contain Gal*p*, highlighted in red. (D) Binding profiles of recombinant hItln1 (50μg/mL) to microbial (left) and mammalian (right) glycan microarrays from NCFG (*12*). Data are shown as mean ± SD (n = 4 technical replicates). (E) Structures of biotin-functionalized carbohydrates used in this study. (F) Schematic of enzyme-linked lectin assay (ELLA). Biotinylated carbohydrates are anchored to surface via streptavidin, and bound lectin is detected using a Strep-HRP antibody and colorimetric HRP substrate. (G) ELLA assay for mItln2 binding to immobilized carbohydrates at varying mItln2 concentrations. Data are shown as mean ± SD (n= 3 technical replicates). (H - J) BLI trace of mItln2 binding to immobilized biotinylated-β-Gal*f* (H), biotinylated-β-Gal*p* (I), and biotinylated-6$SO_3$ LacNAc (J). Binding was tested with 1.5 μM mItln2 for β-Gal*f* and β-Gal*p*, and 5 μM mItln2 for 6$SO_3$ LacNAc. Data were normalized by background subtraction using biotin-loaded streptavidin. (K) BLI response of mItln2 binding to immobilized biotinylated-Gal*p* at 1060 s (depicted in Fig. 2D) was plotted against mItln2 concentration and fitted to a single site binding equation to determine KD. (L and M) Dot blot analysis for different amounts of gastric mucus on nitrocellulose membrane was performed with 0.5 μM mItln2, either in the presence of $Ca^{2+}$ or EDTA. Controls included no lectin (L) and treatment with 10 mM carbohydrates (galactose or glucose) (M). Binding of mItln2 was detected using a Strep-HRP antibody. (N) Spectroscopic assay for crosslinking of 0.05% (w/v) MUC5AC with varying amounts of mItln2 (with and without 100 mM galactose), measured by the increase in absorbance at 405 nm. Data are shown as mean ± SD (n = 3 technical replicates). (O) Time-lapse images of 0.01% (w/v) fluorescently labeled MUC5AC (yellow) without mItln2 (magenta) treatment as a control for experiment depicted in Figure 2H. Binding of mItln2 was detected with Strep antibody. Scale bars, 20 μm. Results in (G), (J), (L), (M), (N), and (O) are representative of two independent experiments. Results in (H), (I), and (K) are representative of three independent experiments. RFU, Relative Fluorescence Unit.

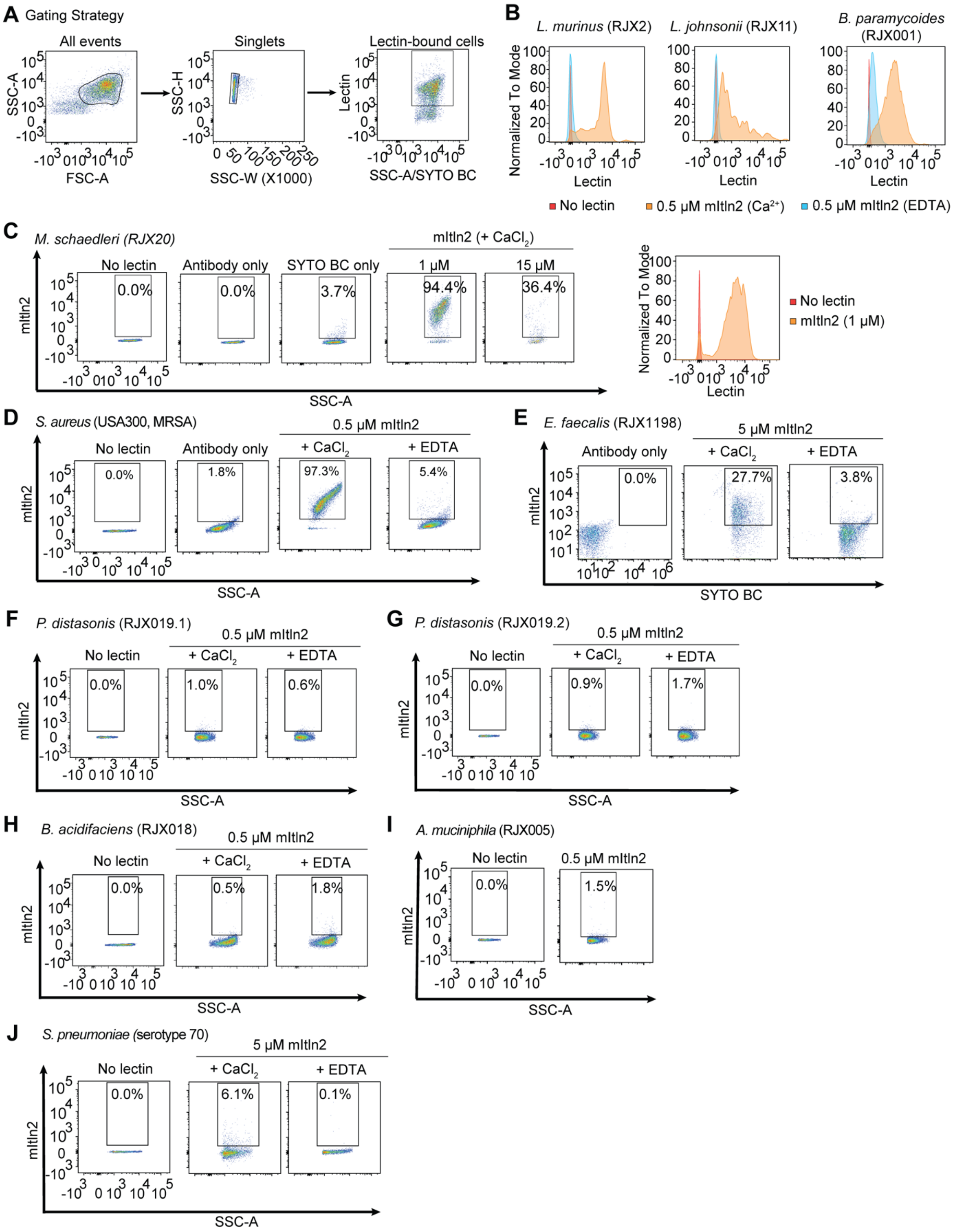
A Gating Strategy
All events
Singlets
Lectin-bound cells
SSC-A
FSC-A
SSC-H
SSC-W (X1000)
Lectin
SSC-A/SYTO BC
B
L. murinus (RJX2)
L. johnsonii (RJX11)
B. paramycoides (RJX001)
Normalized To Mode
Lectin
No lectin
0.5 μM mItln2 (Ca2+)
0.5 μM mItln2 (EDTA)
C
M. schaedleri (RJX20)
mItln2 (+ CaCl2)
No lectin
Antibody only
SYTO BC only
1 μM
15 μM
0.0%
0.0%
3.7%
94.4%
36.4%
mItln2
SSC-A
No lectin
mItln2 (1 μM)
D
S. aureus (USA300, MRSA)
0.5 μM mItln2
No lectin
Antibody only
+ CaCl2
+ EDTA
0.0%
1.8%
97.3%
5.4%
E
E. faecalis (RJX1198)
5 μM mItln2
Antibody only
+ CaCl2
+ EDTA
0.0%
27.7%
3.8%
SYTO BC
F
P. distasonis (RJX019.1)
0.5 μM mItln2
No lectin
+ CaCl2
+ EDTA
0.0%
1.0%
0.6%
G
P. distasonis (RJX019.2)
0.5 μM mItln2
No lectin
+ CaCl2
+ EDTA
0.0%
0.9%
1.7%
H
B. acidifaciens (RJX018)
0.5 μM mItln2
No lectin
+ CaCl2
+ EDTA
0.0%
0.5%
1.8%
I
A. muciniphila (RJX005)
No lectin
0.5 μM mItln2
0.0%
1.5%
J
S. pneumoniae (serotype 70)
5 μM mItln2
No lectin
+ CaCl2
+ EDTA
0.0%
6.1%
0.1%

**Supplementary Fig 3: Binding profiles of mItln2 to different gram-positive and gram-negative microbial isolates.**

(**A**) Flow cytometry gating strategy used to assess lectin binding to microbes.

(**B**) Flow cytometry histogram plots showing mItln2 binding to *L. murinus* RJX2, *L. johnsonii* RJX11, and *B. paramycoides* RJX001 under $Ca^{2+}$ or EDTA conditions. mItln2 was detected by Strep antibody. Unstained samples (no lectin) served as a control. (**C – J**) Flow cytometry analysis of mItln2 binding to *M. schaedleri* RJX020 (C), *S. aureus* MRSA (D), *E. faecalis* RJX1198 (E), *P. distasonis* RJX019.1 (F), *P. distasonis* RJX019.2 (G), *B. acidifaciens* RJX018 (H), *A. muciniphilia* RJX005 (I), and *S. pneumoniae* serotype 70 (J). Dot plots show lectin binding (anti-Strep DY549) vs. SSC or nucleic acid stain (SYTO BC). Histograms display cell counts as a percent of the maximum signal against lectin binding in different conditions. Unstained samples (no lectin) or samples treated with SYTO BC or Strep antibody alone served as controls. Data in (B), (C), (F), (G), (H), (I), and (J) are representative of two independent experiments. Data in (D), (E) are representative of three independent experiments.

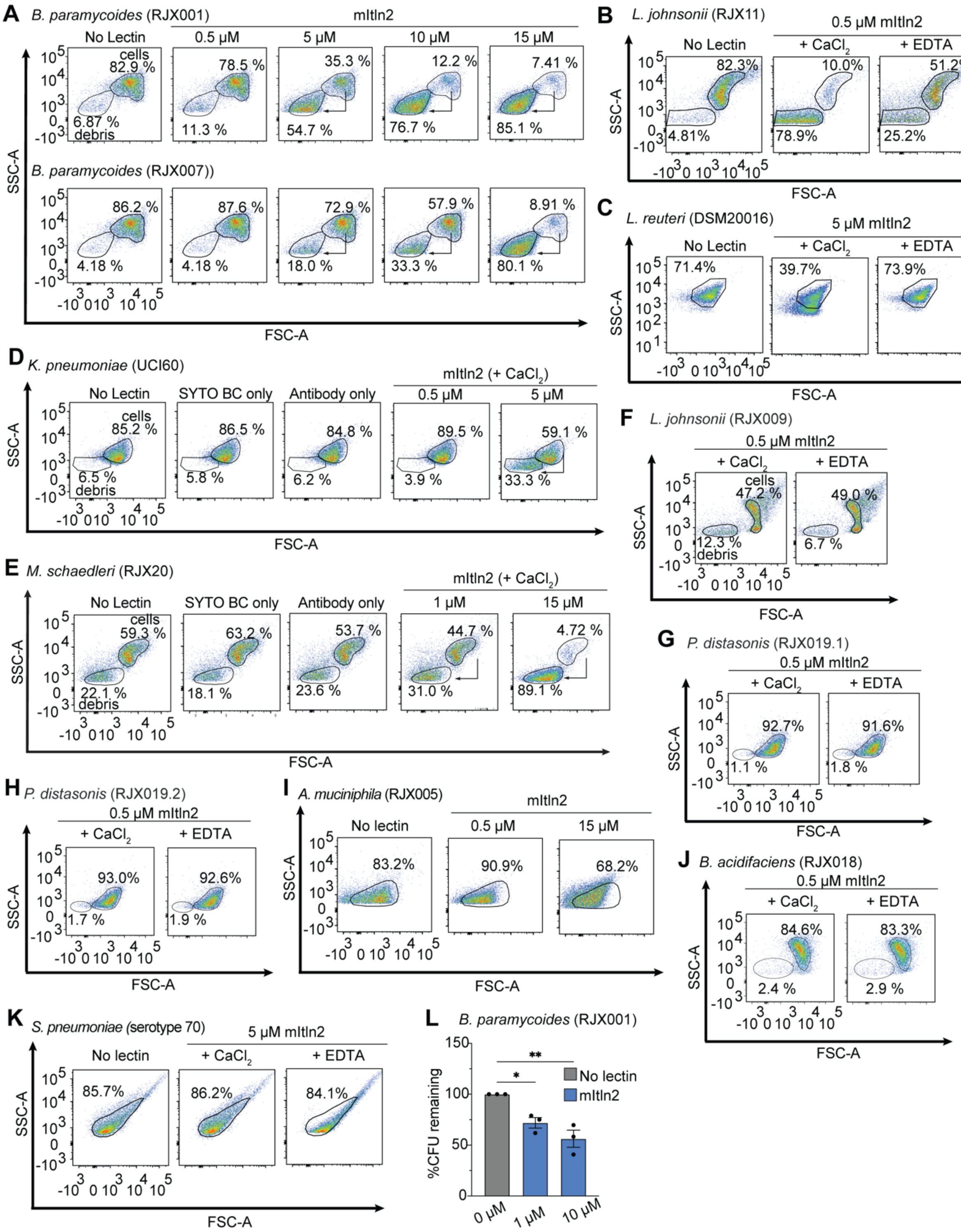

A B. paramycoides (RJX001)
mItln2
No Lectin
0.5 μM
5 μM
10 μM
15 μM
cells
82.9 %
78.5 %
35.3 %
12.2 %
7.41 %
6.87 %
debris
11.3 %
54.7 %
76.7 %
85.1 %
B. paramycoides (RJX007))
86.2 %
87.6 %
72.9 %
57.9 %
8.91 %
4.18 %
4.18 %
18.0 %
33.3 %
80.1 %
SSC-A
FSC-A
B L. johnsonii (RJX11)
0.5 μM mItln2
No Lectin
+ CaCl2
+ EDTA
82.3%
10.0%
51.2%
4.81%
78.9%
25.2%
C L. reuteri (DSM20016)
5 μM mItln2
71.4%
39.7%
73.9%
D K. pneumoniae (UCI60)
mItln2 (+ CaCl2)
SYTO BC only
Antibody only
85.2 %
86.5 %
84.8 %
89.5 %
59.1 %
6.5 %
5.8 %
6.2 %
3.9 %
33.3 %
E M. schaedleri (RJX20)
1 μM
59.3 %
63.2 %
53.7 %
44.7 %
4.72 %
22.1 %
18.1 %
23.6 %
31.0 %
89.1 %
F L. johnsonii (RJX009)
47.2 %
49.0 %
12.3 %
6.7 %
G P. distasonis (RJX019.1)
92.7%
91.6%
1.1 %
1.8 %
H P. distasonis (RJX019.2)
93.0%
92.6%
1.7 %
1.9 %
I A. muciniphila (RJX005)
No lectin
83.2%
90.9%
68.2%
J B. acidifaciens (RJX018)
84.6%
83.3%
2.4 %
2.9 %
K S. pneumoniae (serotype 70)
85.7%
86.2%
84.1%
L B. paramycoides (RJX001)
%CFU remaining
mItln2
0 μM
1 μM
10 μM

**Supplementary Fig. 4: Assessment of cell integrity in mItln2-binding and non-binding microbes.**

(**A** – **K**) Flow cytometry analysis of cell integrity of *B. paramycoides* RJX001 and RJX007 (A), *L. johnsonii* RJX11 (B), *L. reuteri* DSM20016 (C), *K. pneumoniae* UCI60 (D), *M. schaedleri* RJX20 (E), *L. johnonii* RJX009 (F), *P. distasonis* RJX019.1 (G), *P. distasonis* RJX019.2 (H), *A. muciniphilia* RJX005 (I), *B. acidifaciens* RJX018 (J), and *S. pneumoniae* serotype 70 (K) following treatment with mItln2 at different concentrations. Dot plots show SSC vs FSC. Regions representing debris and bacterial cells are indicated. Unstained samples (no lectin) or samples treated with SYTO BC or Strep antibody alone served as controls. Data in (A), (B), (C), (D), (E), (F), (G), (H), (I), (J), and (K) are representative of two independent experiments. (**L**) Quantification of viable *B. paramycoides* RJX001 by dilution plating after incubation with various concentrations of mItln2 for 4 hours. Data show mean ± SD (n = 3 independent experiments). $*P < 0.05$, $**P < 0.01$ (one-way ANOVA followed by Dunnett's multiple comparisons test).

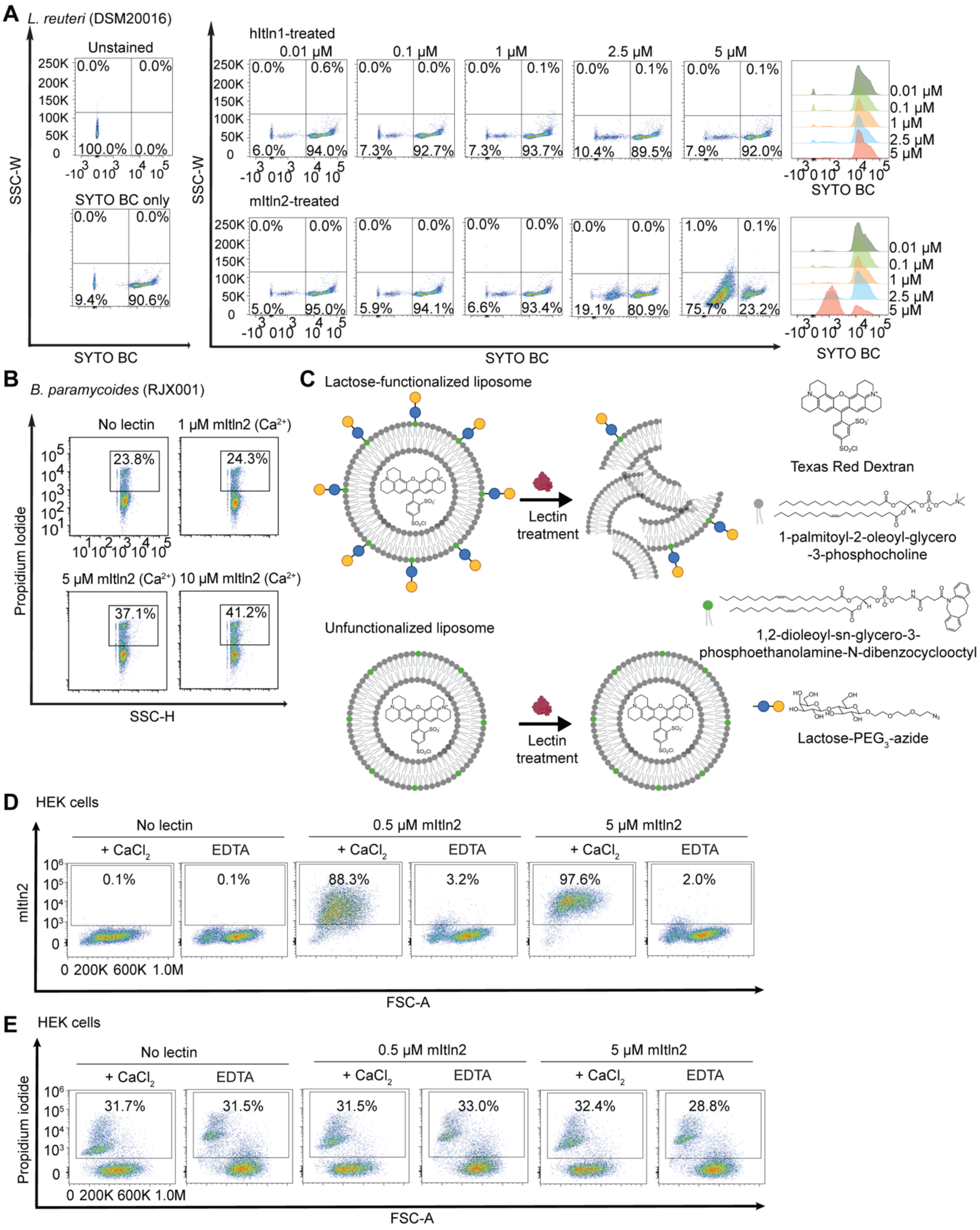
A
L. reuteri (DSM20016)
Unstained
SYTO BC only
hItln1-treated
mItln2-treated
0.01 μM
0.1 μM
1 μM
2.5 μM
5 μM
SSC-W
SYTO BC
B
B. paramycoides (RJX001)
No lectin
1 μM mItln2 (Ca²⁺)
5 μM mItln2 (Ca²⁺)
10 μM mItln2 (Ca²⁺)
23.8%
24.3%
37.1%
41.2%
Propidium Iodide
SSC-H
C
Lactose-functionalized liposome
Unfunctionalized liposome
Lectin treatment
Texas Red Dextran
1-palmitoyl-2-oleoyl-glycero -3-phosphocholine
1,2-dioleoyl-sn-glycero-3- phosphoethanolamine-N-dibenzocyclooctyl
Lactose-PEG3-azide
D
HEK cells
No lectin
0.5 μM mItln2
5 μM mItln2
+ CaCl2
EDTA
0.1%
0.1%
88.3%
3.2%
97.6%
2.0%
mItln2
FSC-A
E
HEK cells
No lectin
0.5 μM mItln2
5 μM mItln2
+ CaCl2
EDTA
31.7%
31.5%
31.5%
33.0%
32.4%
28.8%
Propidium iodide
FSC-A

**Supplementary Fig. 5: Assessment of microbial and mammalian cell viability after mItln2 treatment.**

(**A**) Flow cytometry assessment of cell integrity in *L. reuteri* DSM20016 after treatment with mItln2 at different concentrations, using SYTO BC labeling. Loss of SYTO signal indicates compromised cells. (**B**) Flow cytometry assessment of cell integrity in *B. paramycoides* RJX001 after treatment with mItln2 at different concentrations, using propidium iodide labeling. Gain of propidium iodide signal indicates compromised cells. (**C**) Schematic of liposome disruption assay. Liposomes (200 nm diameter) composed of palmitoyl oleoyl phosphatidylcholine (POPC) with 10 mol% cholesterol for membrane fluidity and 10 mol% 1,2-dioleoyl-sn-glycero-3-phosphoethanolamine-N-dibenzocyclooctyl (18:1 DBCO PE) for glycan attachment were used. The liposomes were encapsulated with Texas red-labeled dextran (m.w. 3000 Da) and functionalized with lactose-$PEG_3$-azide via strain-promoted azido-alkynyl cycloaddition. Addition of mItln2 to the lactose-functionalized liposomes (top) led to a concentration-dependent increase in absorbance at 405 nm and dye release, indicating lipid bilayer disruption. Liposomes without glycan functionalization (bottom) served as controls and showed no membrane disruption, confirming the glycan specificity of the effect. (**D**) Flow cytometry analysis of mItln2 binding to HEK293 cells under $Ca^{2+}$ or EDTA conditions. mItln2 was detected by Strep antibody. Dot plots show lectin binding (anti-Strep DY549) vs. FSC. (**E**) Flow cytometry assessment of cell integrity in HEK293 cells after treatment with mItln2 at different concentrations, using propidium iodide labeling. Gain of propidium iodide signal indicates increased membrane permeability of cells. Data in (A) and (B) are representative of three independent experiments. Data in (D) and (E) are representative of two independent experiments.

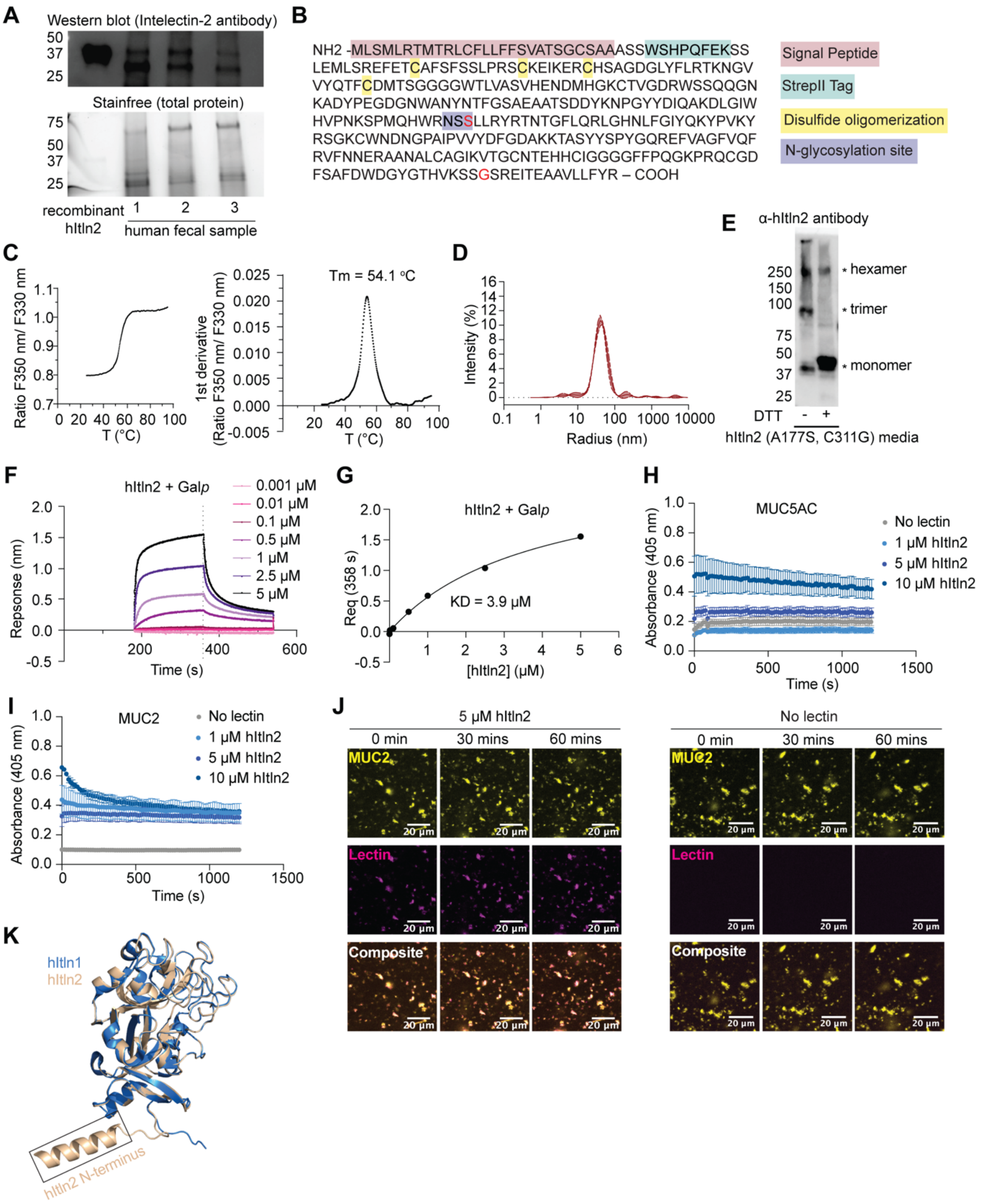
A
Western blot (Intelectin-2 antibody)
Stainfree (total protein)
recombinant hItln2
1 2 3
human fecal sample
B
NH2 -MLSMLRTMTRLCFLLFFSVATSGCSAAASSWSHPQFEKSS
LEMLSREFETCAFSFSSLPRSCKEIKERCHSAGDGLYFLRTKNGV
VYQTFCDMTSGGGGWTLVASVHENDMHGKCTVGDRWSSQQGN
KADYPEGDGNWANYNTFGSAEAATSDDYKNPGYYDIQAKDLGIW
HVPNKSPMQHWRNSSLLRYRTNTGFLQRLGHNLFGIYQKYPVKY
RSGKCWNDNGPAIPVVYDFGDAKKTASYYSPYGQREFVAGFVQF
RVFNNERAANALCAGIKVTGCNTEHHCIGGGGFFPQGKPRQCGD
FSAFDWDGYGTHVKSSGSREITEAAVLLFYR – COOH
Signal Peptide
StrepII Tag
Disulfide oligomerization
N-glycosylation site
C
Ratio F350 nm/ F330 nm
T (°C)
Tm = 54.1 °C
1st derivative (Ratio F350 nm/ F330 nm)
D
Intensity (%)
Radius (nm)
E
α-hItln2 antibody
hexamer
trimer
monomer
DTT
hItln2 (A177S, C311G) media
F
hItln2 + Galp
0.001 μM
0.01 μM
0.1 μM
0.5 μM
1 μM
2.5 μM
5 μM
Repsonse (nm)
Time (s)
G
hItln2 + Galp
KD = 3.9 μM
Req (358 s)
[hItln2] (μM)
H
MUC5AC
No lectin
1 μM hItln2
5 μM hItln2
10 μM hItln2
Absorbance (405 nm)
Time (s)
I
MUC2
No lectin
1 μM hItln2
5 μM hItln2
10 μM hItln2
Absorbance (405 nm)
Time (s)
J
5 μM hItln2
No lectin
0 min
30 mins
60 mins
MUC2
Lectin
Composite
20 μm
K
hItln1
hItln2
hItln2 N-terminus

**Supplementary Fig. 6: Expression and characterization of recombinant hItln2.**
(**A**) Western blot (top) and stain-free gel (bottom) of fecal samples from three individuals under reducing conditions. In Western blot, hItln2 was detected by an intelectin-2 specific antibody. Recombinant StrepII-tagged hItln2 was used as a control. (**B**) Amino acid sequences of StrepII-tagged hItln2, with the signal peptide (pink), StrepII-tag (cyan), cysteine residues (needed for oligomerization, yellow), and N-glycosylation motifs (purple) highlighted. Mutations (A177S and C311G) introduced to optimize recombinant hItln2 expression are labeled in red. (**C**) Ratio of change in intrinsic fluorescence intensity at 330 nm and 350 nm (left) and its first derivative (right) as a function of temperature for recombinant hItln2, obtained from differential scanning fluorimetry. (**D**) Relative distribution of particle sizes for hItln2 protein, as measured by dynamic light scattering. (**E**) Western blot of hItln2 protein under reducing or non-reducing conditions, visualized with pan-intelectin antibody. Molecular weight of monomeric hItln2 is 34 kDa, and different oligomers are indicated by *. (**F**) BLI trace of hItln2 binding to immobilized biotinylated-β-Gal*p* at varying concentrations. Data were normalized by background subtraction, using biotin-loaded streptavidin. (**G**) BLI response of hItln2 binding to immobilized biotinylated-Galp at 358 seconds, plotted against hItln2 concentration and fitted to a single-site binding equation, yielding a KD of 3.9 µM. (**H** and **I**) Spectroscopic assay measuring crosslinking of 0.05% (w/v) MUC5AC (I) and MUC2 (J) by hItln2, indicated by the increase in absorbance at 405 nm. Data are shown as mean ± SD (n = 3 technical replicates). (**J**) Time-lapse images of 0.01% (w/v) fluorescently labeled MUC2 (yellow) treated with 5 µM hItln2 (magenta). Binding of hItln2 was detected with Strep antibody. MUC2 without lectin treatment was used as a control. Scale bars, 20 µm. **(K)** Alignment of hItln1-monomer (blue, PDB ID 4WMY) and hItln2-monomer (wheat, predicted model), highlighting conserved protein structures. The lengthened N-terminal sequence in hItn2, predicted to form an alpha helix, is labeled. Results in (A), (C), (D), and (E) are representative of three independent experiments. Results in (F), (G), (H), (I), and (J) are representative of two independent experiments

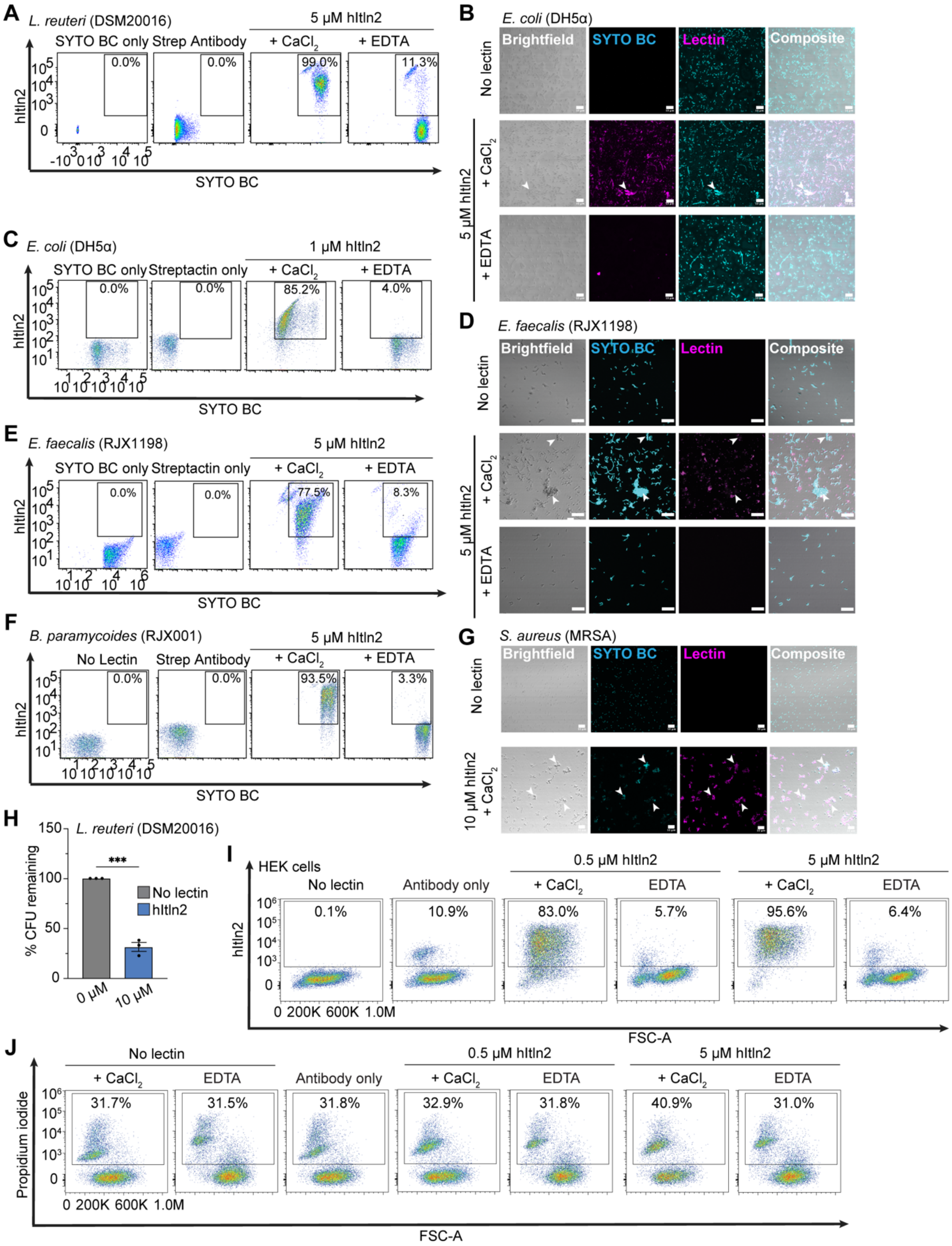
A
L. reuteri (DSM20016)
5 μM hItln2
SYTO BC only
Strep Antibody
+ CaCl2
+ EDTA
0.0%
0.0%
99.0%
11.3%
hItln2
SYTO BC
B
E. coli (DH5α)
Brightfield
SYTO BC
Lectin
Composite
No lectin
5 μM hItln2
+ CaCl2
+ EDTA
C
E. coli (DH5α)
1 μM hItln2
SYTO BC only
Streptactin only
+ CaCl2
+ EDTA
0.0%
0.0%
85.2%
4.0%
hItln2
SYTO BC
D
E. faecalis (RJX1198)
Brightfield
SYTO BC
Lectin
Composite
No lectin
5 μM hItln2
+ CaCl2
+ EDTA
E
E. faecalis (RJX1198)
5 μM hItln2
SYTO BC only
Streptactin only
+ CaCl2
+ EDTA
0.0%
0.0%
77.5%
8.3%
hItln2
SYTO BC
F
B. paramycoides (RJX001)
5 μM hItln2
No Lectin
Strep Antibody
+ CaCl2
+ EDTA
0.0%
0.0%
93.5%
3.3%
hItln2
SYTO BC
G
S. aureus (MRSA)
Brightfield
SYTO BC
Lectin
Composite
No lectin
10 μM hItln2 + CaCl2
H
L. reuteri (DSM20016)
% CFU remaining
***
No lectin
hItln2
0 μM
10 μM
I
HEK cells
0.5 μM hItln2
5 μM hItln2
No lectin
Antibody only
+ CaCl2
EDTA
+ CaCl2
EDTA
0.1%
10.9%
83.0%
5.7%
95.6%
6.4%
hItln2
FSC-A
J
No lectin
0.5 μM hItln2
5 μM hItln2
+ CaCl2
EDTA
Antibody only
+ CaCl2
EDTA
+ CaCl2
EDTA
31.7%
31.5%
31.8%
32.9%
31.8%
40.9%
31.0%
Propidium iodide
FSC-A

**Supplementary Fig. 7: Binding profiles of hItln2 to different microbial and mammalian cells.**

(**A**) Flow cytometry analysis of hItln2 binding to *L. reuteri* DSM20016 under $Ca^{2+}$ or EDTA conditions, detected by StrepII antibody. Dot plots showing lectin binding are displayed as anti-Strep DY549 vs. SYTO BC. (**B** and **C**) Microscopy images (B) and flow cytometry analysis (C) of hItln2 binding to *E. coli* DH5α under $Ca^{2+}$ or EDTA conditions. hItln2 was detected by Streptactin. Scale bars, 10 µm. (**D** and **E**) Microscopy images (D) and flow cytometry analysis (E) of hItln2 binding to *E. faecalis* RJX1198 under $Ca^{2+}$ or EDTA conditions. hItln2 was detected by Streptactin. Scale bars, 20 µm. (**F**) Flow cytometry analysis of hItln2 binding to *B. paramycoides* RJX1001 under $Ca^{2+}$ or EDTA conditions, detected by StrepII antibody. Dot plots showing lectin binding are displayed as anti-Strep DY549 vs. SYTO BC. **(G)** Images of *S. aureus* MRSA stained with hItln2 (magenta) and counterstained with SYTO BC (teal). hItln2 was detected by StrepII antibody. Scale bars, 10 µm. (**H**) Quantification of viable *L. reuteri* DSM20016 by dilution plating after 4-hours incubation with various concentrations of hItln2. Data show mean ± SEM (n = 3 independent experiments). ***P < 0.001 [unpaired two-tailed t-test]. (**I**) Flow cytometry analysis of hItln2 binding to HEK293 cells under $Ca^{2+}$ or EDTA conditions. hItln2 was detected by Strep antibody. Dot plots show lectin binding (anti-Strep DY549) vs. FSC. (**J**) Flow cytometry assessment of cell integrity in HEK293 cells after treatment with hItln2, using propidum iodide labeling. Gain of propidium iodide signal indicates compromised cells. Treatment with SYTO BC and Strep antibody or Streptactin (without lectin) served as a control in (B), (D), and (F). Unstained samples (no lectin) or those treated with SYTO BC, Strep antibody, or Streptactin alone served as controls in (A), (C), (E), (F), (I), and (J). Examples of microbial agglutination are marked with white arrowheads in (B), (D), and (G). Results in (A), (B), (D), (I), (J), and (K) and are representative of two independent experiments. Result in (C), (E), (F), and (G) are representative of three independent experiments.

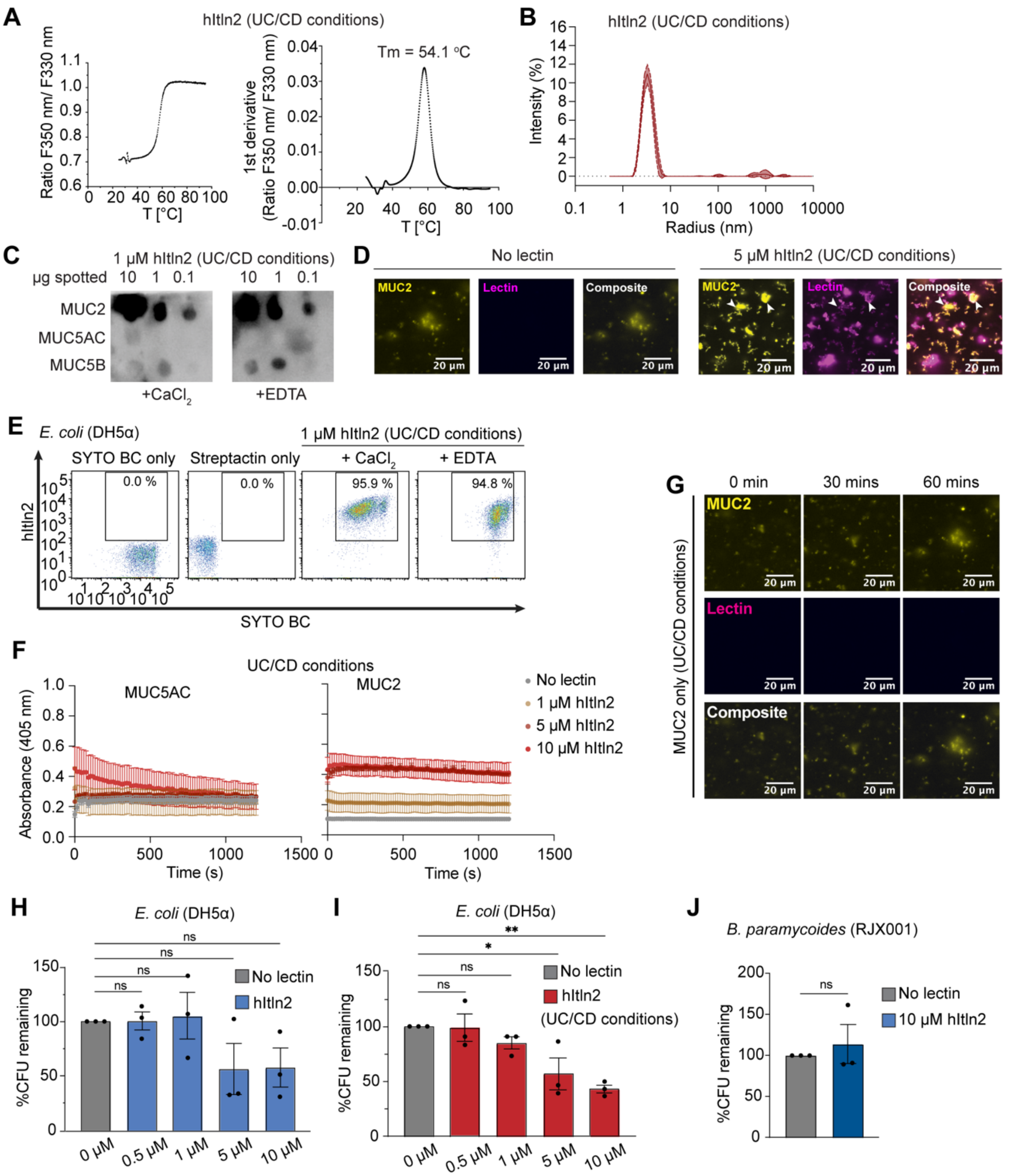

A
hItln2 (UC/CD conditions)
Ratio F350 nm/ F330 nm
T [°C]
Tm = 54.1 °C
1st derivative
(Ratio F350 nm/ F330 nm)
B
hItln2 (UC/CD conditions)
Intensity (%)
Radius (nm)
C
1 μM hItln2 (UC/CD conditions)
μg spotted 10 1 0.1 10 1 0.1
MUC2
MUC5AC
MUC5B
+CaCl2
+EDTA
D
No lectin
5 μM hItln2 (UC/CD conditions)
MUC2
Lectin
Composite
20 μm
E
E. coli (DH5α)
1 μM hItln2 (UC/CD conditions)
SYTO BC only
Streptactin only
+ CaCl2
+ EDTA
0.0 %
0.0 %
95.9 %
94.8 %
hItln2
SYTO BC
F
UC/CD conditions
MUC5AC
MUC2
Absorbance (405 nm)
Time (s)
No lectin
1 μM hItln2
5 μM hItln2
10 μM hItln2
G
0 min
30 mins
60 mins
MUC2 only (UC/CD conditions)
MUC2
Lectin
Composite
20 μm
H
E. coli (DH5α)
%CFU remaining
No lectin
hItln2
ns
0 μM
0.5 μM
1 μM
5 μM
10 μM
I
E. coli (DH5α)
%CFU remaining
No lectin
hItln2
(UC/CD conditions)
ns
*
**
0 μM
0.5 μM
1 μM
5 μM
10 μM
J
B. paramycoides (RJX001)
%CFU remaining
No lectin
10 μM hItln2
ns

**Supplementary Fig. 8: Characterization of hItln2 activity at low pH and low salt (UC/CD conditions).**

(**A**) Ratio of change in intrinsic fluorescence intensity at 330 nm and 350 nm (left) and its first derivative (right) as a function of temperature for recombinant hItln2 at low pH and salt concentration (UC/CD conditions), obtained from differential scanning fluorimetry. (**B**) Relative distribution of particle sizes for hItln2 protein at low pH and salt concentration (UC/CD conditions), analyzed by dynamic light scattering. (**C**) Dot blot analysis of MUC2, MUC5AC, and MUC5B probed with 1 µM hItln2 in $Ca^{2+}$ or EDTA under low pH and salt concentration (UC/CD conditions), detected by Strep-HRP antibody. (**D**) Images of 1 µM hItln2 (magenta) binding to 0.01% (w/v) fluorescently labeled MUC2 (yellow) under low pH and salt concentration (UC/CD conditions), detected by Strep antibody. MUC2 without lectin treatment served as a control. Scale bars, 20 µm. **(E)** Flow cytometry of 1 µM hItln2 (UC/CD) binding to *E. coli* DH5α (E) in $Ca^{2+}$ and EDTA conditions under low pH and salt concentrations (UC/CD conditions), detected by Streptactin. Unstained samples (no lectin) or those treated with SYTO BC, Strep antibody, or Streptactin alone served as controls. (**F**) Spectroscopic assay of 0.05% (w/v) MUC5AC and MUC2 crosslinking by hItln2, measured by the increase in absorbance at 405 nm. Data are shown as mean ± SD (n = 3 technical replicates). (**G**) Time-lapse images of 0.01% (w/v) fluorescently labeled MUC2 (yellow) without any mItln2 (magenta) treatment as a control for experiment depicted in Fig. 5G. Binding of hItln2 was detected with Strep antibody. Scale bars, 20 µm. (**H** and **I**) Quantification of viable *E. coli* DH5α by dilution plating after 4-hours incubation with various concentrations of hItln2 under physiological pH and salt conditions (H) and under low pH and salt concentration (UC/CD conditions) (I). Data show mean ± SEM (n = 3 independent experiments). ***P < 0.001, ns: not significant [one-way ANOVA followed by Dunnett's multiple comparisons test for (H) and (I). (**J**) Quantification of viable *B. paramycoides* RJX001 after 4-hours treatment with 10 µM hItln2. Data show mean ± SEM (n = 3 independent experiments). ns: not significant [unpaired two-tailed t-test]. Results in (A), (B), (C), and (E), are representative of three independent experiments. Result in (D), (F), and (G) are representative of two independent experiments.

| **Read Count (% total)** | **SI (tissue)** | **CTRL (enteroids)** | **IL-4 (enteroids)** | **IL-13 (enteroids)** | **IL-22 (enteroids)** |
|---|---|---|---|---|---|
| Total | 815974 | 1509287 | 1590206 | 1180500 | 1238489 |
| *Itln1* | 773631 (94.8%) | 1464695 (97.0%) | 1102941 (69.4%) | 810944 (68.7%) | 1203291 (97.2%) |
| *Itln2* | 42343 (5.2%) | 44592 (3.0%) | 487265 (30.6%) | 369556 (31.3%) | 35198 (2.8%) |
| *Itln3* | ND | ND | ND | ND | ND |
| *Itln4* | ND | ND | ND | ND | ND |
| *Itln5* | ND | ND | ND | ND | ND |
| *Itln6* | ND | ND | ND | ND | ND |

**Supplementary Table 1.** Read count (mRNA) and relative abundance of intelectin paralogs from small intestine (SI) and enteroids derived from B6.C-Itln1-6 mice, as determined by next-generation sequencing. CTRL = control, ND = not detected.

### METHODS

#### Congenic C57BL/6NTac.BALB/cAnNTac-Itln1:6 (B6.C-Itln1-6) mouse model

All animal experiments were approved by the Institutional Animal Care and Use Committee at the University of California, Davis. Male BALB/cAnNTac mice, encoding a full *Itln1-6* locus[18], were crossed with female C57BL/6NTac mice that encode a single intelectin gene, *Itln1* (parent strains obtained from Taconic Biosciences, Germantown, NY). Male offspring, positive for the *Itln1-6* locus (F1/N1: heterozygous), were crossed with pure C57BL/6NTac female mice for an additional nine generations (N10: ~99.9% C57BL/6NTac background). Experimental tissues and enteroids were derived from heterozygous mice. Animals were humanely euthanized under deep anesthesia with ketamine/xylazine (100/10 mg/kg).

#### Genotyping

DNA was isolated from tissue samples by HotSHOT method[58]. Briefly, samples were incubated in alkaline lysis buffer (25 mM NaOH, 0.2 mM EDTA, pH 12.0) overnight in a water bath at 65 °C. Following digestion, an equivalent volume of neutralization buffer (40 mM Tris-HCl, pH 5.0) was added to the digest solution. The *Itln1-6* haplotype was identified using primers designed to amplify both *Itln1* and *Itln6*, where the *Itln1* allele (i.e., wild-type C57BL/6N; deletion) generates a 305 bp product, and the *Itln6*-containing allele generates an 88 bp product. Genotyping primers: F: 5'-TATTCCTGTCTCAGCTCCTAG-3', and R: 5'-GTCACAGGTAAAKCCAGAAGG-3' (K = G or T). Polymerase chain reaction (PCR) was performed using an Applied Biosystems thermocycler (Waltham, MA). The PCR products were amplified using *Taq* DNA polymerase (New England Biolabs, Cat# M0273S). Thermocycler conditions: one cycle: 95 °C (30 sec), 35 cycles: (95 °C [30 sec], 62 °C [30 sec], and 70 °C [30 sec]), and additional extension for 5 mins at 70 °C.

#### Enteroid culture

Small intestinal crypts were isolated as previously described with minor modification[59]. Briefly, 10 cm of distal small intestine was flushed with Dulbecco's PBS (DPBS), dissected longitudinally, scraped with a glass coverslip to remove villi, and minced into ~1 cm pieces. Intestinal fragments were incubated at 4°C under gentle rocking (60 rpm) for 45 mins in ice-chilled 2 mM EDTA (DPBS) dissociation solution. The intestinal pieces were allowed to settle at the bottom of the conical tube by gravity, and the dissociation solution was decanted. Following a second 45 mins incubation, intestinal pieces were vigorously shaken for ~3 mins in dissociation solution to release intestinal crypts. The resulting crypt-containing suspension was passed through a 70 µm filter to exclude villi fragments, pelleted, and enumerated. The crypts were plated on 24 well Costar® culture dishes (StemCell Technologies, Cat# 38017) as 50 µL suspensions of 50% v/v growth factor reduced matrigel (Corning, Cat# 356231) in murine Intesticult™ Organoid Growth Media (StemCell Technologies, Cat# 06005), supplemented with 50 µg/mL gentamycin (Sigma-Aldrich, Cat# G1397). Enteroids were passaged every 12 days following standard procedures (StemCell Technologies, document# 28223), and complete media was replenished every three days. For cytokine treatment, enteroids were cultured for nine days prior to 72 hr treatment with recombinant (PeproTech, Cranbury, NJ) murine IL-4 (Cat# 214-14, 20 ng/mL), IL-13 (Cat# 210-13, 10 ng/mL), or IL-22 (Cat# 210-22; 20 ng/mL).

#### RNA extraction, cDNA synthesis, and RT-qPCR

RNA extraction from intestinal tissue was performed using the guanidine thiocyanate/cesium chloride gradient method, as previously reported[18,60]. For enteroids, media was removed from the 24-well culture plates following cytokine treatment, and 500 µL of TRIzol™ reagent (ThermoFisher Scientific, Cat#15596026) was added directly to the Matrigel domes. RNA was extracted as outlined by the manufacturer with minor modifications. Following RNA precipitation and decanting of isopropyl alcohol, resulting pellets were resuspended in 250 mL of 70% molecular grade ETOH (30% v/v DEPC $H_2O$) with 10 mL of 3M sodium acetate, and incubated overnight at -80 °C. Thereafter, the RNA was again pelleted by centrifugation, washed with 80% ETOH, and resuspended in DEPC $H_2O$ to determine concentration by ultraviolet absorbance (Nanodrop™) spectroscopy. cDNA synthesis was performed as previously reported, using 1-3 mg of isolated RNA with the SuperScript™ III First-Strand Synthesis kit (ThermoFisher Scientific, Cat# 18080051), followed by column purification using the QIAquick PCR purification kit (Qiagen, Cat# 28104)[18]. RT-qPCR was performed using a Roche Lightcycler® 2.0 under conditions parallel to those previously reported[18,60]. Primers: *Actb* forward 5'-GGCTGTATTCCCCTCCATCG-3', *Actb* reverse 5'-CCAGTTGGTAACAATGCCATGT-3'. *Itln1* forward 5'- ACCGCACCTTCACTGGCTTC-3', *Itln1* reverse 5'- CCAACACTTTCCTTCTCCGTATTTC-3'. *Itln*-common forward 5'-GCCTCAGCAGAGAAAGGTTCC-3', *Itln*-common reverse 5'-GAAGGTCTGGTAGATGACACCATTC-3'. *Clca1* forward 5'-CCTGACTCCTGACTTCTTAGC-3', *Clca1* reverse 5'-TGAACACCTCACTGCTTGG-3'. *Reg3g* forward 5'-CCTCAGGACATCTTGTGTC-3', *Reg3g* reverse 5'-TCCACCTCTGTTGGGTTCA-3'[18,60].

**Next-generation sequencing of Itln paralogs**

A subset of PCR reactions, generated by RT-qPCR using the *Itln*-common primers, were pooled (B6.C-*Itln1*-6: distal small intestine; n = 3 animals, and enteroids; n = 6 independent samples/treatment), column purified (QIAquick PCR purification kit) and submitted for illumina sequencing (GENEWIZ Amplicon-EZ, Azenta Life Sciences). Sequence data (i.e., unique reads) were binned and then aligned to the reference mRNA for the individual *Itlns* (*Itln1*, *Itln2*, *Itln3*, *Itln4*, *Itln5*, and *Itln6*) retrieved from the BAC contig of the 129S7 strain (Gene Bank #HM370554), which like BALB/c (parental donor strain for the B6.C-*Itln1*-6 congenic mouse model), encodes a full *Itln1-6* locus[18]. Percentage total and individual *Itln1-6* read count (Fig. 1c and 1d, SI Table 1) was generated following two selective filters of the raw data: (1) 283 nt (i.e., expected PCR product size), and (2) the minimum recorded unique read count was set at 2000, representing ≤ 0.2% of total reads (median total read count of experimental groups = 1.23 million; Table S1).

**Cloning of StrepII-tagged mouse intelectin-2 and StrepII-tagged human intelectin-2**

Forward primer (5'GCGTTTAAACTTAAGCTTCACCATGACCCAACTGGGCTTCCTG3') and reverse primer (5'CCACCACACTGGACTAGTGGATCCTCATTAGCGATAAAACAGAAGCACAGC3') were used to amplify strep-mItln2 (Accession number AAO60215) from pFastBac-Strep-mItln2 vector (Kiessling lab, unpublished). Forward primer (5'-AGTTAAGCTTCACCATGCTGTCCATGCTGAGGACAATGACC-3') and Reverse primer (GCTCGGATCCTCATTATCTATAGAACAAGAGTACAGCCGCCTCCGTT) were used to amplify hItln2 cDNA (AY358905) (Kiessling Group, unpublished). The resulting PCR products were inserted into pcDNA4 using Gibson ligation. For hItln2, StrepII-tag was inserted using two-step Quikchange mutagenesis with the following primers: (5'GCAGCAGCCTCTTCTtggagccatccgcagtttgaaaagtcttctCTTGAGATGCTCTCG 3') and (5' CGAGAGCATCTCAAGagaagacttttcaaactgcggatggctccaAGAAGAGGCTGCTGC 3'). Plasmid sequences were verified using Sanger sequencing (Quintara Biosciences) with pCMV forward (5'CGCAAATGGGCGGTAGGCGTG3') and BGH reverse primers (5' CTAGAAGGCACAGTCGAGG 3').

**Recombinant protein expression and purification**
Recombinant hItln1, mItln2, and hItln2 with N-terminal Strep-tag II were expressed by transient transfection of suspension-adapted HEK293 cells, following established procedures[12]. HEK293 cells were maintained at a density of $1x10^6$ cells/mL in DMEM medium (ThermoFisher, Cat# 11995) supplemented with 10% heat-inactivated FBS, 1x penicillin-streptomycin, 1x L-glutamine, and 1x nonessential amino acids in spinner flasks at 37 °C, 5% $CO_2$. Cells were passaged every 2-3 days, with the addition of Pluronic-F68 (ThermoFisher). Transfection was carried out at $1x10^6$ cells/mL in the growth medium using Lipofectamine 2000 (ThermoFisher, Cat# 11668030) as per the manufacturer's protocol. Six hours post-transfection, the culture medium was replaced with FreeStyle F17 expression medium (ThermoFisher, Cat# A1383501), supplemented with 1x penicillin-streptomycin, 1x L-glutamine, 1x non-essential amino acids. Transfected cells were cultured for up to 3 days, and the expression medium was harvested by centrifugation and sterile filtration.

To verify protein expression, expression media was combined with 6x Laemmli buffer containing dithiothreitol, boiled at 95 °C for 10 mins, and run on 4-15% TGX gel (BioRad). Proteins were transferred to PVDF membrane, blocked in 5% milk in TBST (Tris-Buffered Saline with Tween 20), and blotted for the presence of mItln2 using 1:5000 rabbit anti-Itln2 (Proteintech, Cat# 11770-1-AP) and goat-anti-rabbit HRP (1:10,000, Jackson, Cat# AB_2313567). hItln2 was detected on protein expression blots using hItln2 specific antibody (Bevins Lab, 1:5000) and goat-anti-rabbit HRP (1:10,000, Jackson, Cat# AB_2313567)[19].

Purification of StrepII-tagged lectins followed established procedures[12]. The harvested expression medium was subjected to avidin (IBA, Cat# 2-0204-015) treatment, resulting in a final concentration of 0.084 mg/mL, as per the IBA protocol. Protein was captured on Strep-Tactin Superflow High-Capacity resin (IBA, Cat# 2-1208-002), which was preequilibrated with HEPES/Ca buffer (20 mM HEPES, 10 mM $CaCl_2$, 150 mM NaCl, pH 7.4). Subsequently, the resin was washed twice with HEPES/EDTA buffer (20 mM HEPES, 1 mM EDTA, 150 mM NaCl, pH 7.4). The StrepII-tagged lectins were then eluted with 5 mM d-desthiobiotin (Sigma-Aldrich) in HEPES/EDTA buffer and concentrated using 10,000-molecular weight cutoff (MWCO) Vivaspin 6 centrifugal filter (Sartorius).

All proteins underwent buffer exchange to HEPES/EDTA for storage except for StrepII-hItln2 (UC/CD conditions). Purification of hItln2 in low pH and low salt (UC/CD condition) was identical, except that the purified protein was buffer exchanged into PIPES/EDTA buffer (10 mM PIPES, 25 mM NaCl, 1 mM EDTA, pH 5.5). Protein concentrations were determined by absorbance at 280 nM, with extinction coefficients and molecular weights calculated for the monomeric form of each protein (without the signal peptide) using the Protparam tool. StrepII-tagged hItln1 had ε = 79925 $cm^{-1}m^{-1}$ and an estimated molecular mass of 34224 Da (monomer). StrepII-tagged mItln2 had ε = 69830 $cm^{-1}m^{-1}$ and an estimated molecular mass of 34212 Da. StrepII-tagged hItln2 had ε = 69955 $cm^{-1}m^{-1}$ and an estimated molecular mass of 34436 Da (monomer).

**Detection of hItln2 in human fecal samples**
Human fecal samples from three independent donors were kindly provided by the Alm lab (MIT)[13]. Fecal samples were supplied as homogenates in glycerol and stored at -80 °C. To evaluate the secretion of hItln2 in the intestinal lumen, human fecal homogenates were scraped from frozen glycerol stock and thawed on ice. The homogenate was centrifuged at 100 x g for 1 min to pellet fibrous material. The supernatant was transferred to a new tube and pelleted at 3000 x g, 5 mins. The pellet was washed twice with PBS, and then masses of pellets were measured. Pellets were suspended in PBS to achieve a concentration of 40 mg/mL. 20 mL of each sample was mixed with 6x Laemmli buffer containing DTT (Dithiothreitol), boiled at 95 °C, and run on 4-15% TGX Stain-Free gel (Bio-Rad). Total protein was visualized using stain-free settings on ChemiDoc XP (BioRad). Proteins were transferred to PVDF, blocked in 5% milk in TBST, and blotted for hItln2 protein using an anti-hItln2 antibody (Bevins Group, 1:5000) and Goat-anti-rabbit HRP (1:10,000, Jackson, Cat# AB_2313567)[19].

**Treatment of mItln2 with PNGase F**
To examine N-glycosylation, 20 µg of recombinant mItln2 was subjected to PNGase F (NEB, Cat# P0704S) treatment according to the manufacturer's recommendations. Following glycosidase treatment, samples were mixed with 6x Laemmli buffer containing dithiothreitol (DTT), boiled at 95 °C, and run on 4-15% TGX stain-free gel (Bio-Rad) with molecular weight standards (Bio-Rad). Gel was imaged on ChemiDoc MP Imaging system using stain-free imaging settings.

**Circular dichroism (CD) analysis**
Recombinant mItln2 was purified for CD analysis as described above, except all washes, elution, and buffer exchanges were performed with PBS buffer. CD spectra from 180 nm to 260 nm with 1 nm wavelength steps were collected on a Jasco H-1500 (Jasco Inc) at 25 °C. Measurements were conducted with 500 µg/mL of mItln2 (in PBS buffer) using a 1 mm cuvette, with PBS buffer serving as the blank. Averaging time was 5 s, settling time was 0.33 s, and readings were taken in triplicate. Thermal denaturation steps were run from 25 °C to 95 °C with 5 °C temperature steps. Scans were averaged, corrected by subtracting the blank, and plotted in GraphPad Prism to calculate melting temperature and visualize spectra. The calculated mean molar ellipticity (θ) is plotted as a function of wavelength (nm) for the protein.

**Differential Scanning Fluorometry (DSF) and Dynamic Light Scattering (DLS)**
Particle light scattering and change in intrinsic fluorescence intensity of recombinant lectins at 330 nm and 350 nM in response to temperature changes were measured using a NanoTemper Prometheus NT.48 instrument. Capillaries were filled with ~10 µL of recombinant lectins (in HEPES/EDTA buffer or PIPES/EDTA buffer), placed into the sample holder, and the temperature was increased from 25 °C to 90 °C. The fluorescence intensity ratio at 350 nm and 330 nm was plotted as a function of temperature, and its first derivative was calculated using the manufacturer's software. Melting temperature was calculated as the inflection point of the ratio

curve. DLS was measured at 25 °C prior to melting temperature analysis. The hydrodynamic radii were determined using buffer-only capillaries as a baseline. The DLS and DSF experiments were conducted using two replicates for each sample.

**Chemical cross-linking of proteins**

For crosslinking of mItln2 protein, 1 mg/mL of mItln2 (in HEPES/EDTA buffer) was combined with BS3 (ThermoFisher, Cat# 21580) crosslinker in HEPES/EDTA buffer to achieve different final concentrations (0.1 mM, 0.25 mM, 0.5 mM, 1.25 mM, 2.5 mM, and 5 mM). Crosslinking was performed at RT for 30 mins. Subsequently, protein mixtures were denatured by adding an SDS loading buffer with DTT. Samples were then heated at 95 °C for 5 mins, separated by SDS-PAGE, and imaged with ChemiDoc MP Imaging system using SYPRO Ruby protein stain or TGX Stain Free technology.

**Enzyme-linked lectin assay (ELLA)**

To assay mItln2 binding to different carbohydrates, a 96-well MaxiSorp ELISA plate (ThermoFisher, Cat# 442404) was coated with 0.5 μg of streptavidin (Agilent, Cat# SA26) in PBS per well. Subsequently, the wells were loaded with 5 μM of biotinylated carbohydrates (in PBS) for 1 hr at RT. After blocking with 5% BSA in HEPES/Ca buffer, the plate was incubated with various concentrations of recombinant lectins in HEPES/Ca/BSA/T buffer (20 mM HEPES, 10 mM $CaCl_2$, 150 mM NaCl, 0.1% BSA, 0.1% Tween-20) for 2 hrs at RT. Following washes with the HEPES/Ca/BSA/T buffer, the wells were treated with anti-StrepMAB-Classic HRP conjugate antibody (1:10000, IBA, Cat#2-1509-001) in HEPES/Ca/BSA/T buffer for 2 hrs at RT. Then, the plates were washed with HEPES/Ca/BSA/T buffer, and the bound StrepII-tagged lectins were detected colorimetrically using 1-Step Ultra TMB-ELISA (ThermoFisher). The reaction was quenched by adding an equal volume of 2M sulfuric acid. Plates were read at 450 nm on Molecular Devices SpectraMax5. Data were analyzed using GraphPad Prism.

**Mucin dot blot**

Purified porcine MUC2, MUC5AC, and MUC5B were generously provided by the Ribbeck Lab at MIT. Partially purified gastric mucus was purchased from Sigma (Cat# M1778). Mucins were hydrated overnight in deionized water, 4 °C, with rotation to achieve a stock concentration of 10 mg/mL. Mucins and gastric mucus were serially diluted in 20 mM HEPES (pH 7.4) and 150 mM NaCl buffer to generate working stocks for downstream assays. For individual mucins, 1 μg of hydrated mucins were spotted onto nitrocellulose and allowed to dry. After blocking with 5% BSA in TBS-T for 1 hr at RT, 0.5 - 1 μM recombinant lectins in HEPES/Ca/BSA/T buffer or HEPES/EDTA/BSA/T buffer (20 mM HEPES, 1 mM EDTA, 150 mM NaCl, 0.1% BSA, 0.1% Tween-20) were applied to blot overnight at 4 °C. To test the glycan specificity of lectin-mucin interactions, the blots were incubated with the lectins in combination with 10 mM galactose in HEPES/Ca/BSA/T buffer. For UC/CD conditions, the blot was incubated with hItln2 in PIPES/Ca/BSA/T buffer (10 mM PIPES, pH 5.5, 25 mM NaCl, 10 mM $CaCl_2$, 0.1% BSA, 0.1% Tween-20) or PIPES/EDTA/BSA/T buffer (10 mM PIPES, pH 5.5, 25 mM NaCl, 1 mM EDTA, 0.1% BSA, 0.1% Tween-20). Blots were washed in TBS-T and treated with anti-StrepMAB-Classic HRP conjugate antibody (1:10000, IBA, Cat# 2-1509-001) in the appropriate buffer. Subsequently, the blot was washed three times with TBS-T and developed in the ChemiDoc MP Imaging system using ECL Prime reagent (Amersham).

For gastric mucus binding, serial dilutions of mucus were spotted on nitrocellulose and blocked as described above. Membranes were incubated with 0.5 μM strepII-mItln2 in HEPES/Ca/BSA/T alone or in combination with 10 mM glucose or 10 mM galactose. Blots were incubated for 1 hr at RT or overnight at 4 °C, processed as described above, and imaged using autoradiography film (Cole-Palmer) and developing reagents (Kodak).

**Mucin and glycopolymer agglutination assay**

To test the agglutination effect of mItln2, we adapted an assay from Järvå et al.[36] 1% (w/v) mucins or glycopolymer were first diluted to 0.35% (w/v) in HEPES/Ca or PIPES/ Ca buffer. 1 μL mucin or glycopolymer was aliquoted into each well of a 384-well plate (black/clear bottom). Using a multichannel pipette, 34 μL of increasing concentrations of Itln2 in HEPES/Ca or PIPES/ Ca buffer or buffer alone was added simultaneously to wells and briefly mixed to achieve a final concentration of 0.01% (w/v) mucin. The plate was immediately monitored for changes in absorbance at 405 nm (A405nm) using Molecular Devices SpectraMax5 via a kinetic assay, taking reads every 10 seconds for 30 mins at RT. Triplicates were averaged and plotted in GraphPad Prism. Data is displayed as changes in A405nm as a function of time.

**Imaging of lectin-mediated mucin agglutination**

Purified porcine MUC2, MUC5AC, and MUC5B were hydrated in deionized water overnight at 4 °C, with rotation to prepare stock concentration of 10 mg/mL. Mucins were labeled with AlexaFluor 568 NHS ester (ThermoFisher, Cat# A20006) as per the manufacturer's recommendations. Labeled mucins were pelleted at 10,000 x g for 10 mins and washed thrice with excess PBS to remove unreacted fluorophore. Mucins were resuspended to a working concentration of 1% (w/v) in HEPES/Ca or PIPES/Ca buffer. Mucins were plated in a lysine-coated 96-well imaging plate, which was followed by the addition of solutions containing 5 μM lectin, StrepMAB-Classic DY649 (IBA, Cat# 2-1569-050, 1:250) in HEPES/Ca/BSA/T or PIPES/Ca/BSA/T. Mucins treated with only buffers were used as a control. The final mucin concentration in the assay was 0.01% (w/v). Following treatment with lectins, the plate was centrifuged briefly at 1000 x g for 1 min and immediately imaged on Molecular Devices IXM HC confocal microscope. Images were acquired at the start of the experiment and every 15 mins after that for a total of 2 hours. Images were analyzed in Fiji, where settings required for the brightest images were applied to all samples. Signal appears lower for untreated mucins for this reason.

**Synthesis of biotin-carbohydrates and glycopolymers**

Synthesis of biotinylated carbohydrates (β-Gal*f*-biotin and β-Gal*p*-biotin) and 60% lactose-functionalized trans-poly(norbornene) polymer, which were synthesized in-house, has been described previously[12,39]. Other biotinylated carbohydrates (α-Gal*p*, β-LacNAc, β-GlcNAc, α-Neu5Ac, β-6-$SO_3$ LacNAc) were purchased from GlycoTech.

**Biolayer interferometry (BLI)**

Lectin binding to biotinylated carbohydrates was assessed using the OctetRed BLI instrument (ForteBio). Biotinylated carbohydrate was loaded onto streptavidin biosensor (ForteBio) for 120 s as a 5 μM solution in PBS buffer. The sensor was subsequently washed in PBS for 60 s, and a baseline was established in HEPES/Ca/BSA/T or PIPES/Ca/BSA/T buffer for 120 s. Various concentration of recombinant lectins (in HEPES/Ca/BSA/T or PIPES/Ca/BSA/T buffer) was then associated for 600 s followed by dissociation in HEPES/Ca/BSA/T or PIPES/Ca/BSA/T buffer for 600 s. The shake rate was maintained at 1000 rpm throughout the experiment, and the binding assays were performed at 30 ºC. Data were normalized through background subtraction using the response from biotin-loaded streptavidin. The results were analyzed using GraphPad Prism.

To determine the dissociation constant for lectin with biotinylated carbohydrates, we conducted the BLI experiments with various lectin concentrations using the same protocol as above. The response at equilibrium was plotted against the concentration of lectin, and the resulting curve was fit to a one-site total non-linear regression equation to determine the equilibrium dissociation constant (GraphPad Prism).

**Assay of Itln2 on microbial and mammalian glycan array**
Lectin binding to mammalian glycans was assessed using the Consortium for Functional Glycomics (CFG) version 5.5 microarray and RayBiotech slides (Cat# GA-Glycan-300-1). Binding to microbial glycans was assessed using the CFG microbial glycan microarray version 2 (MGM). Each glycan sample on the microarray had six replicates. The microarray slides were first rehydrated for 5 mins in HEPES/Ca/BSA/T buffer. Subsequently, the slides were incubated with 25 µg/mL of StrepII-tagged mItln2 in HEPES/Ca/BSA/T buffer for 1 hr at RT. The slides were then washed with HEPES/Ca/BSA/T buffer and incubated with StrepMAB-Classic DY-549 (1:250, IBA, Cat# 2-1566-050) in HEPES/Ca/BSA/T buffer for 1 hr at RT. Afterward, the array went through washing steps with HEPES/Ca/BSA/T buffer, HEPES/Ca buffer, and deionized water. Finally, the bound lectin in the microarray was detected using a fluorescent scanner. The results are presented as relative fluorescence units obtained by averaging the background-subtracted signals for the four replicate spots (after excluding the highest and lowest values from the six replicates), with error bars representing the SD of the averaged values.

**Assay for lectin binding to fecal samples**
Freshly collected murine fecal samples were homogenized in PBS. Human fecal homogenates were scraped from frozen glycerol stock and thawed on ice. The homogenate was centrifuged at 100 x g for 1 min to pellet fibrous material. The supernatant was transferred to a new tube and pelleted at 3000 x g, 5 mins. The pellet was washed twice with PBS, and the optical density at 600 nm ($OD_{600}$) was measured. To test Itln2 binding to fecal microbiota, the homogenate was diluted to $OD_{600}$ of 0.2 and treated with 0.6 µM of StrepII-tagged mItln2, StrepMAB-Classic DY549 antibody (1:250, IBA, Cat# 2-1566-050), or 1 µM StrepII-hItln2, StreptactinXT-DY549 (1:250, IBA, Cat# 2-1565-050) and SYTO BC (1:1000, ThermoFisher, Cat# S34855) in HEPES/Ca/BSA/T buffer for 2 hrs at 4 ºC. To test $Ca^{2+}$-dependent binding of Itln2, staining was carried out in HEPES/EDTA/BSA/T buffer and used as a control. For the hIltn2 in UC/CD condition, the homogenate was treated with 1 µM StrepII-hItln2, StreptactinXT-DY549 (1:250, IBA, Cat# 2-1565-050) and SYTO BC (1:1000, ThermoFisher, Cat# S34855) in PIPES/Ca/BSA/T buffer or PIPES/EDTA/BSA/T buffer for 2 hrs at 4 ºC. Untreated fecal samples and the fecal samples treated solely with StrepMAB-Classic DY549 antibody, StrepTactinXT-549, or SYTO BC were used as additional controls. Following staining, the cells were analyzed on LSRII, LSR Fortessa HTS flow cytometer or FACS Symphony flow cytometer. Flow data were analyzed with FlowJo.

For the microscopy experiment, the fecal samples were processed identically to the flow experiment described earlier. Following staining, the samples were transferred to a lysine-coated 96-well black/clear bottom Plate (ThermoFisher, Cat# 165305) and centrifuged at 500 x g for 1 min. Imaging was performed using the RPI Spinning Disk Confocal microscope or Molecular Devices IXM HC confocal microscope. Image analysis was performed using Fiji.

**Assay for lectin binding to microbial isolates**
Bacterial isolates *Bacillus paramycoides* (RJX001), *Bacillus paramycoides* (RJX007), *L. reuteri* (RJX004), *L. murinus* (RJX002), and *L. johnsonii* (RJX009) were isolated from mouse stool by Xavier Lab (Broad Institute). *Lactobacillus reuteri* (DSM 20016) and *Escherichia coli* DH5α were obtained from Prof. Federico Rey, courtesy of Dr. Robert Kerby (UW-Madison) and NEB, respectively. *B. paramycoides* (RJX001), *B. paramycoides* (RJX007), *L. reuteri* (DSM 20016 and RJX004), *L. murinus* (RJX002), and *L. johnsonii* (RJX009) were grown anaerobically in a chamber with AnaeroGen sachet (ThermoFisher) at 37 °C for 16 to 24 hrs without shaking. *B. paramycoides* (RJX001 and RJX007) were grown in CHG medium [brain heart infusion ($37gL^{-1}$, BD) supplemented with 1% vitamin K1-hemin (ATCC), D-(+)-cellobiose ($1gL^{-1}$, SigmaAldrich), D-(+)-fructose ($1gL^{-1}$, SigmaAldrich), D-(+)-maltose ($1gL^{-1}$, SigmaAldrich), and L-(+)-cysteine ($1gL^{-1}$, SigmaAldrich). *L. reuteri* (DSM 20016 and RJX004), *L. murinus* (RJX002), and *L. johnsonii* (RJX009) were grown in MRS medium (BD). *E. coli* was grown in Luria Broth (LB) at 37 °C for 16 to 24 hrs with shaking at 200 rpm.

To test lectin binding, cells were revived from glycerol stock in their respective buffers and grown overnight in appropriate aerobic or anaerobic conditions. Cultures were harvested by centrifugation and washed with PBS twice before $OD_{600}$ measurement. Staining was performed at $OD_{600}$ of 0.2 for all samples. For staining, cells were treated with various concentrations of StrepII-tagged lectins, StrepMAB-Classic DY549 antibody (1:250, IBA, Cat# 2-1566-050) or StreptactinXT-DY549 (1:250, IBA, Cat# 2-1565-050), and SYTO BC (1:1000, ThermoFisher, Cat# S34855) in HEPES/Ca/BSA/T buffer for 2 hrs at 4 °C. To test the $Ca^{2+}$-dependent binding of lectins, staining was carried out in HEPES/EDTA/BSA/T buffer and used as a control. For UC/CD conditions, cells were treated with lectins in PIPES/Ca/BSA/T or PIPES/EDTA/BSA/T buffers. Unstained bacteria, bacteria treated only with StrepMAB-Classic DY549 antibody or StreptactinXT-DY549, and bacteria treated only with SYTO BC were used as additional controls. Following staining, the cells were diluted five-fold and analyzed on LSR Fortessa HTS flow cytometer or FACS Symphony flow cytometer. Data were analyzed with FlowJo.

For the microscopy experiment, the microbial isolates were processed identically to the flow experiment described earlier. Following staining, the samples were transferred to a lysine-coated 96-well black/clear bottom Plate (ThermoFisher, Cat# 165305) and centrifuged at 500 x g for 1 min. Imaging was performed using Molecular Devices ImageXpress Micro Confocal Microscope, RPI spinning disc confocal, or Olympus FV1200 Laser Scanning Confocal Microscope. Image analysis was performed using Fiji.

**Assay for lectin binding to pathogenic isolates *(S. aureus, K. pneumoniae,* and *S. pneumoniae*)**

*Staphylococcus aureus* USA300Lac (MRSA) and *Klebsiella pneumoniae* UCI60 were generously shared by Xavier Lab (Broad Institute) and Hung Lab (Broad Institute), respectively. Cells were grown in suspension of LB at 37 $^{o}$C for 16 to 24 hrs with shaking at 200 rpm.
*S. pneumoniae* (Klein) Chester serotypes 70 (ATCC10370) were obtained from ATCC. Cells were grown in suspension of Todd Hewitt broth (BD) with 0.5% yeast (BD) without shaking at 37 $^{o}$C under 5% CO2 for 24 hrs.

To test lectin binding, cells were harvested by centrifugation, washed with PBS, and fixed in 4% formaldehyde in PBS for 1 hr at RT. Fixation was neutralized with Tris buffer (pH 7.4) at a working concentration of 50 mM. After washing cells with PBS, $OD_{600}$ was measured, and staining was performed at $OD_{600}$ of 0.2. For staining, fixed cells were treated with various concentrations of StrepII-tagged lectins, StrepMAB-Classic DY549 antibody (1:250, IBA, Cat# 2-1566-050) or StreptactinXT-DY549 (1:250, IBA, Cat# 2-1565-050), and SYTO BC (1:1000, ThermoFisher, Cat# S34855) in HEPES/Ca/BSA/T buffer for 2 hrs at 4 $^{o}$C. To test $Ca^{2+}$-dependent binding of lectins, staining was carried out in HEPES/EDTA/BSA/T buffer and used as a control. For UC/CD conditions, cells were treated in PIPES/Ca/BSA/T or PIPES/EDTA/BSA/T buffers. Unstained bacteria, bacteria treated only with StrepMAB-Classic DY549 antibody or StreptactinXT-DY549, and bacteria treated only with SYTO BC were used as additional controls. Following staining, the cells were diluted five times without washing and analyzed on LSR Fortessa HTS flow cytometer or FACS Symphony flow cytometer. Data were analyzed with FlowJo.

For the microscopy experiment, the microbes were processed identically to the flow experiment as described earlier. Following staining, the fixed samples were transferred to a lysine-coated 96-well black/clear bottom Plate (ThermoFisher, Cat# 165305) and centrifuged at 500 x g for 1 min. Imaging was performed using Molecular Devices ImageXpress Micro Confocal Microscope or Olympus FV1200 Laser Scanning Confocal Microscope. Image analysis was performed using Fiji.

**Colony forming unit (CFU) Assays**

*L. reuteri* (DSM 20016) was cultured anaerobically in MRS medium at 37 $^{o}$C for 16 hrs without shaking. *B. paramycoides* RJX001 grown anaerobically in a chamber with AnaeroGen sachet (ThermoFisher) at 37 $^{o}$C for 16 to 24 hrs without shaking. *S. aureus* USA300Lac (MRSA) was cultured in LB medium at 37 $^{o}$C for 16 hrs with shaking at 200 rpm. Upon reaching the mid-logarithmic phase, the cells were harvested by centrifugation and washed with PBS before measuring $OD_{600}$. Subsequently, cells were diluted to $OD_{600}$ of 0.2 and treated with various concentrations of StrepII-tagged lectins in HEPES/Ca/BSA/T or PIPES/Ca/BSA/T buffer for 4 hrs (at 4 $^{o}$C for *L. reuteri* and *S. aureus* and 37 $^{o}$C for *B. paramycoides*). Following incubation with lectins, cells were serially diluted in sterile HEPES/Ca ($10^{1}$-$10^{5}$) or PIPES/Ca buffer and plated on appropriate agar plates (MRS for *L. reuteri*, CHG for *B. paramycoides*, or LB for *S. aureus*) and incubated overnight at 37 $^{o}$C. *L. reuteri* and *B. paramycoides* plates were grown in Oxoid AnaeroJar 2.5 L anaerobic chamber with AnaeroGen 2.5 L sachet (ThermoFisher, AN0025A). Surviving bacterial colonies were counted, and CFU/mL was calculated. Data is displayed as a percentage relative to the control sample (cells treated with HEPES/Ca/BSA/T buffer or PIPES/Ca/BSA/T buffer without lectins).

**Suspension-culture growth assay for lectin-treated microbes**
For lectin-mediated growth inhibition assays, *K. pneumoniae* UCI60 was cultured directly from glycerol stock in LB media overnight at 37 °C with shaking. Cells were back-diluted in fresh LB and grown to mid-log phase ($OD_{600}$ of 0.4 – 0.7). Cells were harvested by centrifugation, washed with PBS, and the $OD_{600}$ was measured. Cells were then diluted in M9 growth media (1x M9 salts, 0.1% glucose, 100 µM $CaCl_2$, 2 mM $MgSO_4$) to achieve a 2x stock of cells ($OD_{600}$ = 0.001). 2x cells were mixed with 2x stocks of increasing concentration of lectins in M9 media in a final volume of 50 µL. The plate was covered with BreathEasy gas-permeable film (Diversified Biotech, BEM-1) and allowed to grow overnight, 18 hrs, at 37 C in SpectraMax M5 plate reader. $OD_{600}$ nm reads were taken every 15 mins. Triplicate samples were averaged, and the mean $OD_{600}$ nm was plotted as a function of time (GraphPad Prism).

As *Lactobacilli* do not grow in M9 minimal media, an alternate growth recovery assay was designed to assay lectin-mediated growth assay. *L. reuteri* (DSM 20016 and RJX004), *L. murinus* (RJX002), and *L. johnsonii* (RJX009) were cultured anaerobically in MRS medium at 37 °C for 16 hrs without shaking. Cells were harvested by centrifugation, washed with PBS, and $OD_{600}$ measured. Subsequently, the cells were diluted to $OD_{600}$ of 0.2 and treated with various concentrations of StrepII-tagged lectins in HEPES/Ca/BSA/T buffer in a sterile 96-well-round bottom plate. Following treatment, cells were diluted five-fold in MRS broth, sealed with film, and allowed to recover overnight on SpectraMax M5 plate reader, 37 °C without shaking, taking $OD_{600}$ reads every 15 mins. The growth assay was run in triplicate, and the data was averaged. Data was displayed as mean $OD_{600}$ changes plotted as a function of time.

**Time-lapse microscopy of L. reuteri**
*L. reuteri* (DSM 20016) was cultured anaerobically in MRS medium at 37 °C for 16 hrs without shaking. To assess the impact of lectin on *L. reuteri*, the bacterial cells were harvested by centrifugation and washed with PBS before $OD_{600}$ measurement. Subsequently, the cells were diluted to $OD_{600}$ of 0.2 and treated with various concentrations of StrepII-tagged lectins in HEPES/Ca/BSA/T buffer in a lysine-coated 96-well black/clear bottom Plate (ThermoFisher, Cat# 165305). Imaging was performed using a 40x water immersion objective in Molecular Devices ImageXpress Micro Confocal Microscope at 10 mins intervals over 6 hours. Images were processed in Fiji.

**Assay for propidium iodide uptake in *B. paramycoides***
*B. paramycoides* RJX001, grown anaerobically in a chamber with AnaeroGen sachet (ThermoFisher) at 37 °C for 16 to 24 hrs without shaking, was harvested at mid-logarithmic phase by centrifugation and washed with PBS before measuring $OD_{600}$. Subsequently, cells were diluted to $OD_{600}$ of 0.2 and treated with various concentrations of StrepII-tagged mItln2 in HEPES/Ca/BSA/T or HEPES/EDTA/BSA/T buffer for 4 hrs at 37 °C. Following incubation with lectins, cells were treated with propidium iodide (ThermoFisher, Cat# P3566) for 20 mins and analyzed using FACS Symphony flow cytometer.

**Assay for lectin binding and propidium iodide uptake in mammalian cells**
HEK-293T cells were maintained in DMEM medium (ThermoFisher, Cat# 11995) supplemented with 10% heat-inactivated FBS, 1x penicillin-streptomycin, 1x L-glutamine, and 1x nonessential amino acids. To assay lectin binding, the HEK-293T cells were dissociated with EDTA and washed with DPBS. Subsequently, cells (100K/ staining condition) were washed with HEPES/Ca or HEPES/EDTA buffer and then incubated with various concentrations of StrepII-tagged lectins and StrepMAB-Classic DY-549 (1:250, IBA, Cat# 2-1566-050) in HEPES/Ca/BSA/T or HEPES/EDTA/BSA/buffer for 3 hrs at 4 ºC. Following incubation with lectins, the cells were treated with 1µL of Propidium iodide (ThermoFisher, Cat# P3566) for 20 mins at 4 ºC. The cells were then washed with the respective buffer twice and analyzed using an Attune flow cytometer

**Liposome preparation**
Palmitoyl oleoyl phosphatidylcholine (POPC) was obtained as a 25 mg/mL solution in $CHCl_3$ from Avanti Polar Lipids (Cat# 850457C). 1,2-dioleoyl-sn-glycero-3-phosphoethanolamine-N-dibenzocyclooctyl (18:1 DBCO PE) was obtained as a 10 mg/mL solution in $CHCl_3$ from Avanti Polar Lipids (Cat# 870129C). The lipid solutions were stored at -20 ºC and used within 3 months. Cholesterol was obtained as a white solid from Sigma Aldrich (Cat# C8667).

Prior to preparing a lipid film, the lipid solutions were warmed to ambient temperature for at least 30 mins to prevent condensation from contaminating the solution and degrading the lipid film. 330 µL of POPC, 75 µL of 18:1 DBCO PE, and 100 µL of a 4 mg/mL cholesterol solution in $CHCl_3$ were added to a glass scintillation vial. The solvent was removed with a gentle stream of nitrogen, and the resulting lipid film was stored under a high vacuum for a minimum of twelve hours prior to use.

The dried lipid film was rehydrated with 1 mL of 150 mM NaCl, 20 mM HEPES, 10 mM $CaCl_2$, pH 7.4 containing 200 ug/mL Texas Red-dextran conjugate MW 3000 Da (Thermo Fisher, Cat# D3329), and vortexed vigorously for approximately 3 mins to form a suspension of multilamellar vesicles (MLVs). To obtain a sufficient quantity of large unilamellar vesicles (LUVs), at least three independent lipid film preparations were pooled together for the subsequent formation of LUVs. The resulting lipid suspension was pulled into a Hamilton (Reno, NV) 1 mL gastight syringe and the syringe was placed in an Avanti Polar Lipids Mini-Extruder (610000). The lipid solution was then passed through a 200 mm pore size hydrophilic polycarbonate filter (Avanti Polar 610006) 21 times, the newly LUV suspension being collected in the syringe that did not contain the original suspension of MLVs to prevent the carryover of MLVs into the LUV solution. The newly formed LUVs were purified via size-exclusion chromatography (SEC) using Sephadex G-50 (Sigma Aldrich G50150) and functionalized with 5 mM β-Lactose-PEG3-azide (Sigma Aldrich SMB00404-25MG) with shaking at 4 °C overnight via a strain-promoted azido-alkynyl cycloaddition (SPAAC). The lactose-functionalized liposomes were then purified by SEC to remove unreacted glycan ligands.

**Liposome membrane disruption assay**
Liposomes were diluted to OD 0.1 in 150 mM KCl, 20 mM HEPES, 10 mM $CaCl_2$, pH 7.4 buffer. Absorbance (405 nm) was measured with a SpectraMax microplate reader (Molecular Devices). 50 µL of the liposome suspension was added to a 96-well plate, and initial

measurement was recorded. Then, 25 µL of mItln2 or vehicle (buffer only) was added to each well, and absorbance was continuously read for 45 mins, with measurements taken every 30 seconds.
For the microscopy experiment to visualize liposomal structural integrity, following measurement of absorbance over time following lectin addition, samples were imaged on an RPI Spinning Disc Confocal. Image analysis was performed using Fiji.

**Approval for human sample research**
Human stool samples used in the flow cytometry test of human intelectin-2 were obtained under a protocol approved by the Massachusetts Institute of Technology [Institutional Review Board (IRB) protocol ID no. 1510271631]. The participants provided informed consent, and all experiments adhered to the regulations of the review boards.

**Statistical analysis**
Statistical analysis of the data was performed using GraphPad Prism 9. Unpaired Student's t-test was used when comparing two independent samples. One-way ANOVA with Dunnett's multiple comparisons test was used when comparing three or more samples. Three biological replicates were used, except when indicated otherwise. ns: not significant, *$p<0.05$, **$p< 0.01$, ***$p<0.001$, ****$p<0.0001$.

**Software**
Protein structures and overlays were generated with PyMol (The PyMol Molecular Graphics System, Version 3.0 Schrodinger, LLC). Homology models of mItln2 were generated with SwissModel[61]. Putative structures of hItln2 were generated with AlphaFold3[62]. Flow cytometry data was analyzed using FlowJo (FlowJo™ v10.8 for Mac Softwarejm). Imaging data was analyzed using Fiji[63]. GraphPad Prism version 10.1.1 for Mac was used for data analysis and graph generation (GraphPad Software, Boston, MA USA, www.graphpad.com). Sequencing data was analyzed using SnapGene (www.snapgene.com). Carbohydrate structures were generated using ChemDraw version 20.1 for Mac (Revvity Signals Software). References were cited using Zotero version 6.0.37 (www.zotero.org).

**Data availability**
The glycan array data for mItln2 and hItln2 are provided in the SI Table S2 and S3. Any additional information required to reanalyze the data reported in this paper is available from the lead contact upon request.
